# Immediate methane and carbon dioxide release from exposed permafrost at an active retrogressive thaw slump in the Canadian Arctic

**DOI:** 10.64898/2026.06.17.732964

**Authors:** Lexi Joyce, Laura L. Lapham, Roger MacLeod, Marcus R. Phillips, Jalal Norooz Oliaee, Adam W. Gillespie, Peter Morse, Scott Dallimore, Jackie Goordial

## Abstract

The Arctic is warming rapidly, causing permafrost thaw and accelerating the release of greenhouse gases. Rapid thaw features such as retrogressive thaw slumps are increasing in frequency and severity across the Arctic; however, their associated greenhouse gas emissions are poorly constrained. Current estimates of emissions from retrogressive thaw slumps rely largely on laboratory incubations and carbon stock estimates rather than *in-situ* field measurements. Here we directly quantify methane and carbon dioxide fluxes from the exposed headwall of an active retrogressive thaw slump. We show that thaw immediately releases biogenic methane and carbon dioxide, originating from gases trapped within the frozen soil matrix. Microbial transcription of methyl-coenzyme M reductase suggests archaea carrying out methanogenesis at subzero temperatures are the source of trapped methane. Carbon emissions varied by an order of magnitude among cryostratigraphic units, reflecting differences in geomorphologic history, organic carbon and nitrogen content, and microbial community composition. Carbon emissions were highest from organic-rich paleo cryosols from the Late Holocene that contained abundant methanogenic archaea. We estimate that ∼300 kg C (CO_2_ equivalents) is emitted annually from the headwall of this small thaw slump (surface area of ∼1200 m^2^). Considering the thousands of active slumps and extensive coastal permafrost erosion across the northern continuous permafrost zone, such features may represent a growing natural source of GHG emissions. These findings indicate that current permafrost carbon feedback models underestimate GHG release by omitting the direct release of trapped gases stored in permafrost.

## Introduction

The Arctic is warming up to four times faster than the rest of the planet, causing permafrost (perennially frozen ground) to thaw^1^. The northern circumpolar permafrost region contains nearly half of the belowground organic carbon on Earth^2,3^. Cryogenic (freeze-thaw) processes in permafrost-affected soils develops cryosols that sequester and accumulate carbon in permafrost over millennia. When permafrost thaws, previously frozen organic matter in these paleo cryosols becomes available for microbial decomposition, producing carbon dioxide (CO_2_) and methane (CH_4_), and potentially transforming this long-term carbon reservoir into a net carbon source. This process creates a positive permafrost carbon feedback (PCF) that can accelerate atmospheric warming^3,4^.

Microbiota are the primary drivers of carbon emissions from thawing permafrost soils. As permafrost thaws, enhanced nutrient accessibility and faster metabolic rates can increase microbially mediated greenhouse gas (GHG) emissions^5^. Decades of permafrost research has focused on GHG production after soil has thawed above 0°C, for example in laboratory incubations that simulate warming, or field studies that directly measure GHG flux from thawed permafrost *in situ*. As a result of the emphasis on post-thaw microbial mineralisation, alternative emission sources have been overlooked in field studies and climate models.

One such pathway includes the direct release of GHGs physically trapped within the frozen soil matrix. Permafrost is known to contain occluded gases such as CH_4_ and CO_2_,^6–11^, yet their potential contributions to emissions upon thaw remains poorly constrained and rarely quantified. Most permafrost incubation methodologies require permafrost to be thawed (>0°C) during experiment setup, inadvertently releasing trapped GHGs and as a result underestimating the true quantity of gases associated with permafrost thaw. The limited studies that have not thawed permafrost prior to warming experiments, or that explicitly measured trapped gases, have all confirmed the presence of occluded gases^8,9,11^. Field-based carbon flux estimates through eddy covariance towers or soil chamber-based measurements cannot differentiate between direct emissions from trapped GHGs or secondary emissions from microbial respiration stimulated by increased temperatures. Due to these difficulties, the direct release of trapped GHGs from permafrost thaw has been modeled^12^ but has never been explicitly measured in field studies.

Most Arctic permafrost is insulated by a seasonally thawed active layer that, together with snow and vegetation, acts as a thermal buffer between permafrost and the atmosphere. The active layer supports diverse methanotrophic microbiota^13^ that can oxidize CH_4_ produced in the thawing lower active layer and permafrost before it reaches the atmosphere. In some instances, high rates of CH_4_ oxidation in the active layer produce net CH_4_ sinks^14,15^. Arctic permafrost currently warms at a rate of up to about 1°C per decade^16^, gradually increasing active layer thickness over decades to centuries^17,18^. In contrast, approximately 3.6 million km^2^ of the northern permafrost region is vulnerable to rapid permafrost thaw^19^ that leads to large ecosystem changes over months to years^18^. Abrupt permafrost thaw is estimated to release up to 80 ± 19 Pg of carbon by the year 2300 under the high emissions Representative Concentration Pathway 8.5 global warming scenario^19^, with CH_4_ comprising ∼20% of emissions^19^. Features such as retrogressive thaw slumps (RTS) initiate when permafrost thaws in ice-rich hillslope terrain^20^ and are increasing in frequency across the Arctic^21–26^. In the Western Canadian Arctic alone, there was a 247% increase in the number of active RTS between 1960 and 2004^22^. In ice-rich permafrost the stability of the ground is supported by ground ice. When the ground ice melts, that support is removed and unconsolidated permafrost materials may collapse into a debris flow. The removal of material exposes a headwall of permafrost that lacks a buffering active layer. Currently, only three studies directly measure GHG emissions from RTS^27–29^ and all focus specifically on secondary emissions from thawed outflow materials rather than primary emissions at the headwall where trapped gas release would be expected. Further, permafrost erosion at 65% of Arctic coasts directly exposes about 6.6 × 10^4^ linear kilometres of permafrost^30^, where the scientific focus on GHG flux has largely overlooked GHGs occluded in permafrost.

Here, we directly measure CH_4_ and CO_2_ fluxes from the exposed permafrost headwall of an active RTS on Niglintgak Island (69°21’26.6"N, 135°20’24.7"W) in the Inuvialuit Settlement Region of the Northwest Territories (Fig. 1). This site is situated in the outer Mackenzie River delta, a region characterised by both biogenic and thermogenic CH_4_ hotspots^31^ and is within the limits of the Laurentide Ice Sheet for northwestern Canada^32^. This RTS is comprised of distinct cryostratigraphic layers that enable us to investigate how geologic history, physicochemical soil characteristics, and microbial community composition may influence GHG flux. We quantify an abundance of methanogenic archaea in soils associated with net CH_4_ emissions, including identifying RNA transcripts of the methyl-coenzyme M reductase, a key gene required for the production of CH_4_. Our results provide the first *in situ* field evidence that RTS headwalls can act as immediate point sources of CH₄ and CO₂, a finding that would extend to coastal permafrost erosional headwalls, and as a result reveal a missing flux pathway that should be incorporated into permafrost carbon models to avoid systematic underestimation of near-term Arctic GHG emissions.

**Fig 1.**
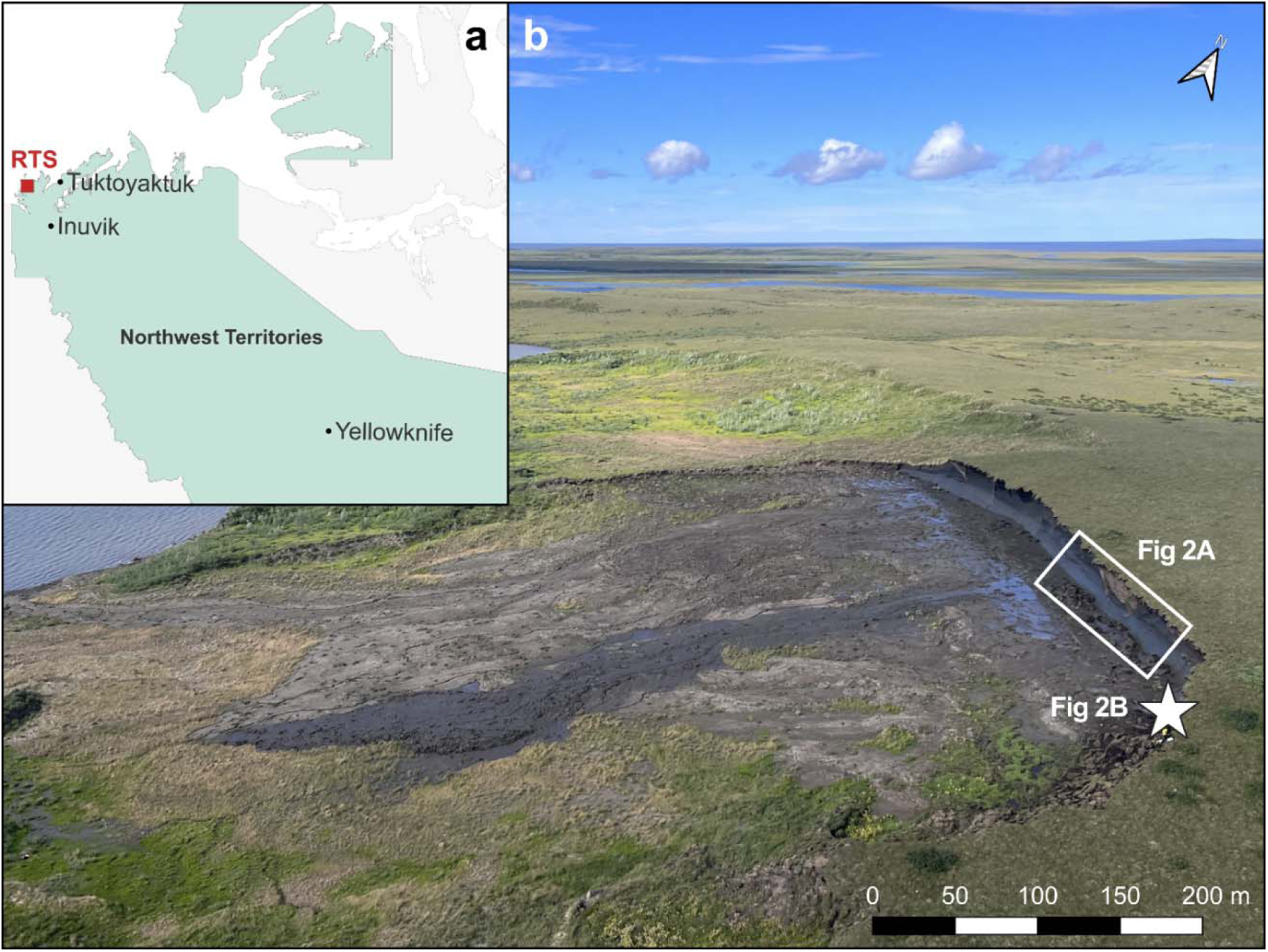
A map indicating the location (red square) of the retrogressive thaw slump (RTS) at Niglintgak Island (69°21’26.6"N, 135°20’24.7"W) in the Northwest Territories, Canada (a) and an aerial photo of the RTS taken in August 2022 (b). In b, the star indicates where flux measurements were taken on the headwall. This RTS is polycyclic and was active in the past but then stabilized as indicated by the vegetated scar in the background, but is presently active as indicated by exposed, ice-rich permafrost and transport of muddy materials down gradient from the foot of the headwall.

## Results

### Characterisation of permafrost headwall cryostratigraphy

The exposed permafrost headwall had four visually distinct cryostratigraphic layers (Units 1 - 4) beneath a modern active layer, together with large ice-wedge inclusions truncated at the active layer base (Fig. 2). Units were identified by a combination of cryostratigraphic observations, physicochemical properties (Table S1), and radiocarbon dating (Table S2).

**Fig 2.**
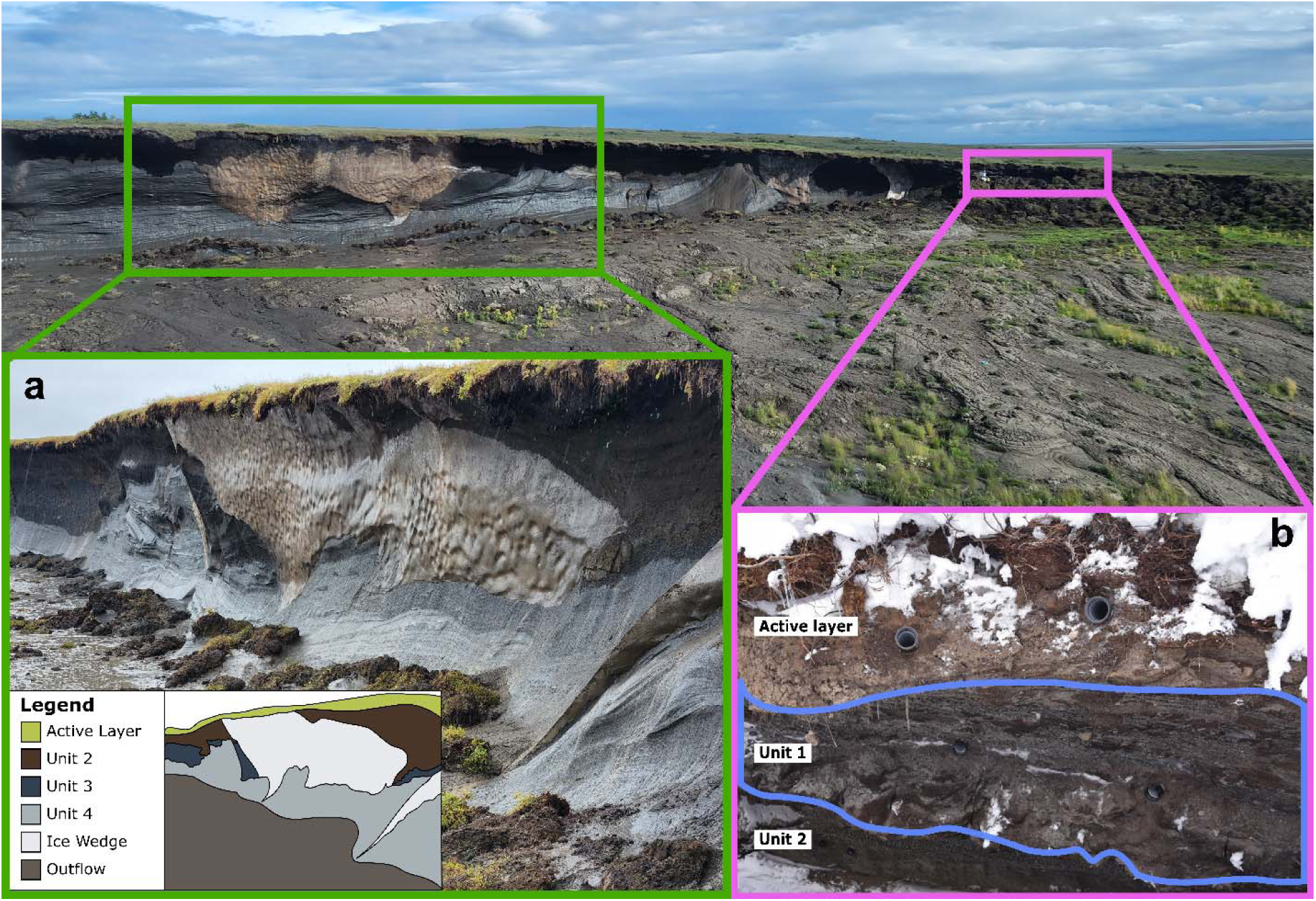
Cryostratigraphy of exposed permafrost at the headwall of the RTS at Niglintgak Island. The West side of the RTS includes the active layer, Unit 2, Unit 3, Unit 4, ice wedge, and thaw outflow which are shown as green, dark brown, blue-grey, light grey, white, and light brown, respectively on the inset (a). Unit 1 (highlighted in blue) is only present on the East side of the RTS (b).

Unit 1 occurs directly beneath the 30–50 cm active layer and is restricted to the eastern/southern flank of the slump where the thaw face intersects a slope with earth hummocks at the surface. It is highly heterogenous, reflecting mixing and burial of soil materials by freeze-thaw and slope processes^33,34^. Unit 1 consists of a heterogeneous mix of dark, organic-rich sandy silt and medium-brown, mineral-rich sandy silt. This unit has variable ice content with thin ice lenses and veins. Bulk sediment and organic detritus in this paleo cryosol dates range from 6180 – 4420 calBP (5260 ± 20 to 4020 ± 20 ^14^C years BP).

Unit 2 is a more homogenous, medium-brown sandy silt with visible abundant roots and a sharp thaw unconformity at its base. This unit has variable ice content with thin ice lenses and veins. Radiocarbon dating yielded an age of 9400 - 9030 calBP (8240 ± 25 ^14^C years BP). We interpret Unit 2 as a paleo-active layer, which is widespread in this region^35^, that developed during the Holocene Thermal Maximum. The unit showed no deformation or cryoturbation.

Units 3 is medium-grey, silty diamicton lacking roots, and is separated from the underlying Unit 4 by a distinct thaw unconformity. A reticulate ice cryostructure indicates slow refreezing of saturated sediments^36^, suggesting that a thaw event occurred after deposition of Unit 4 followed by refreezing and incorporation into permafrost of supraglacial melt-out till. Radiocarbon ages are (40 380 – 39 070 calBP; 34400 ± 290 ^14^C years BP).

Unit 4 is the basal unit and is comprised of deformed layers of massive ice and icy beds with medium-grey silt rich diamicton. The folded and deformed sediment is interpreted to result from glaciotechtonic deformation with no evidence of thaw. Radiocarbon dating of Unit 4 yielded a date range of 41 440 – 40 080 calBP (35 800 ± 340 ^14^C years BP), which is relatively consistent with the non-finite date (>37 000 ^14^C BP) reported for the same locality^32^.

Across Units 2 through 4 there is no evidence of cryoturbation, indicating stratigraphic preservation since deposition. Radiocarbon dates for Units 3 and 4 predate the Late Wisconsinan advance of the Laurentide Ice Sheet that began 27 000 – 30 000 ^14^C years BP and overrode the region by about 23 000 – 30 000 ^14^C years BP^37^. Thus, we do not interpret these radiocarbon dates as depositional ages because the presence of re-worked organic material in these units, including much older wood and coal, make a minimum date difficult to assign. A detailed geologic and paleoenvironmental interpretation is included in the Supplementary Materials.

### Greenhouse gases are released immediately upon permafrost thaw and are microbial in origin

Greenhouse gas fluxes measured using soil chambers installed at the headwall of the RTS revealed substantial emissions of CH_4_ and CO_2_ released directly from the permafrost exposure. Gas fluxes ranged by an order of magnitude across cryostratigraphic units (Fig. 3). Methane fluxes ranged from 4.8 x 10^-4^ to 5.0 g CH_4_-C m^-2^ day^-1^ and CO_2_ fluxes ranged from 1.4 x10^-3^ to 6.1 g CO_2_-C m^-2^ day^-1^. The highest GHG fluxes were from Unit 1, although fluxes within this unit were highly variable. While most fluxes measured in Unit 1 were of a similar magnitude to those in the other cryostratigraphic units, several wall chambers (samples 1-2, 1-4, and 1-10 for CH_4_ flux; samples 1-2 and 1-10 for CO_2_) had fluxes that were much higher than the rest of the samples (Fig. 3; Table S3). As a result, average CH_4_ fluxes were significantly higher in Unit 1 than in Units 2 and 4 and CO_2_ fluxes were significantly higher in Unit 2 than Unit 3 (Kruskal-Wallis and Dunn, *p* < 0.001).

**Fig 3.**
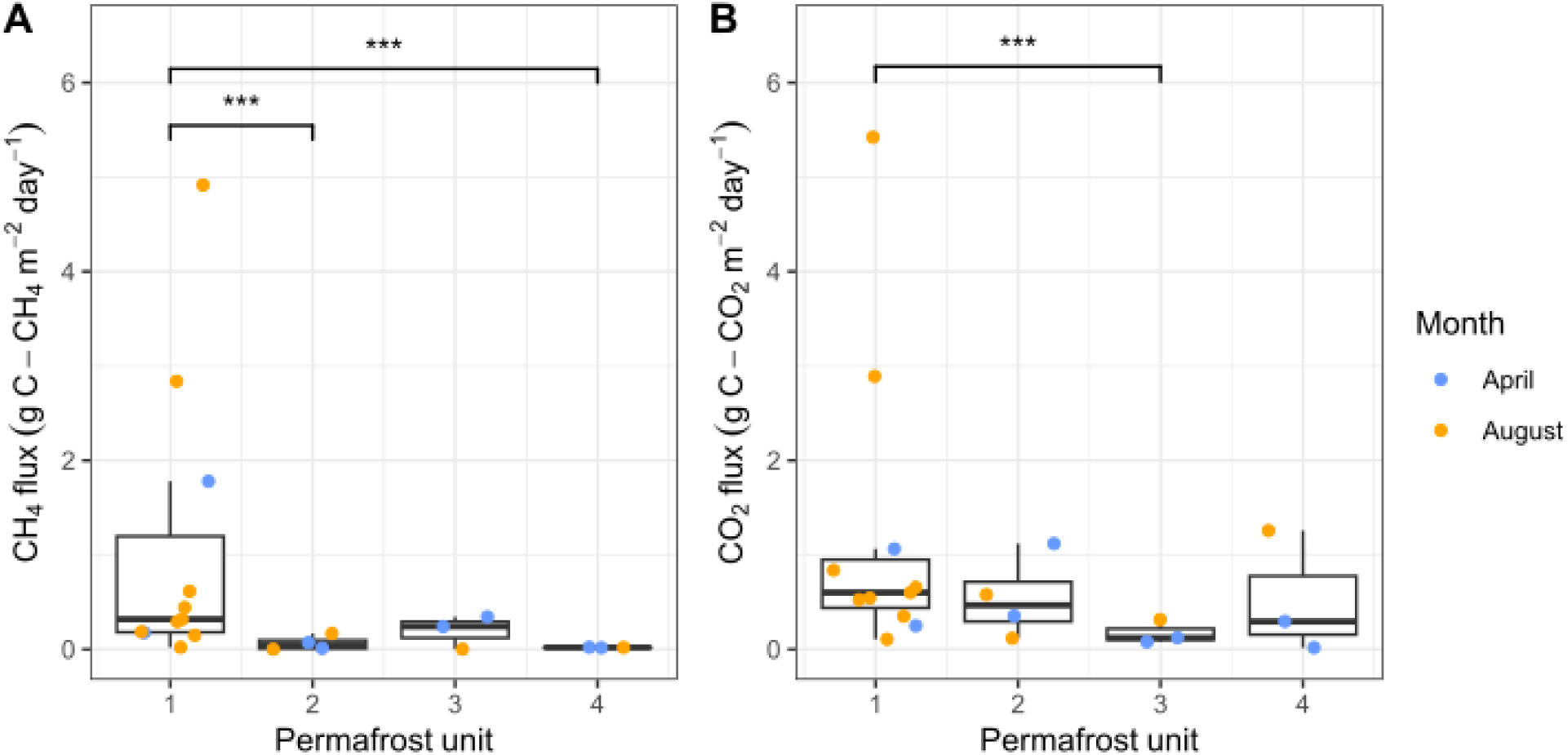
In-situ CH_4_ (a) and CO_2_ (b) fluxes, measured from the RTS headwall immediately as permafrost thawed. Triplicate flux measurements were taken at each wall chamber, and each point represents the mean measurement from each wall chamber. No outliers were removed to account for the heterogeneity within a single permafrost unit. Each boxplot is defined by the first and third quartile values, with the middle line representing the median. The whiskers extend from the lowest value within 1.5 IQR of the first quartile to the highest value within 1.5 IQR of the third quartile. Difference in fluxes between permafrost units was determined by performing the Kruskal-Wallis test and Dunn’s post-hoc test with a Holm-Bonferroni adjustment (*** *p* < 0.001).

The variation in GHG flux across cryostratigraphic layers reflected underlying physicochemical differences (Table S1). CH_4_ emissions from all wall chambers were positively correlated with total organic carbon (TOC; Spearman’s *rho* = 0.65, *p* < 0.001) and total nitrogen (TN; Spearman’s *rho* = 0.68, *p* < 0.001), suggesting nutrient availability modulates CH_4_ release. Unit 1 had significantly higher and heterogeneous TOC (2.29 to 39.15%) and TN (0.52 to 1.69%) than the other cryostratigraphic units (Kruskal-Wallis and Dunn, *p* < 0.001). Gravimetric water content was positively correlated with CH_4_ emissions (Spearman’s *rho* = 0.61, *p* < 0.001). In contrast, CO_2_ fluxes did not correlate with any measured physicochemical properties, indicating that CO_2_ emissions may be shaped by processes other than bulk abiotic characteristics of the permafrost.

Methane and CO_2_ emissions did not differ significantly between summer and winter measurements (Fig 3; Kruskal-Wallis and Dunn, *p* > 0.05), supporting that trapped gases were the source of GHG emissions. Consistent with that interpretation, occluded gases were extracted in the laboratory and were confirmed to be present in all permafrost cores. Trapped CH_4_ ranged from 1.2 x 10^-5^ – 1.6 x 10^-1^ mg C-CH_4_ g^-1^ and CO_2_ ranged from 1.3 x 10^-2^ – 8.3 x 10^-1^ mg C-CO_2_ g^-1^ permafrost (Table S1). These values represent minimum trapped gas estimates, as gas loss likely occurred because of abrasive drilling, transport and handling, and the increase in diffusive surface area following sample extraction (see Supplemental for trapped gas methods).

To determine the origin of the emitted gases, we measured stable carbon isotope ratios (δ^13^C-CH_4_; δ^13^C-CO_2_) and concentrations of CH_4_ and CO_2_ from gases collected at the end of each flux measurement period in 2022 (Fig. 4, Table S4). The Unit 3 sample was lost in transit.

**Fig 4.**
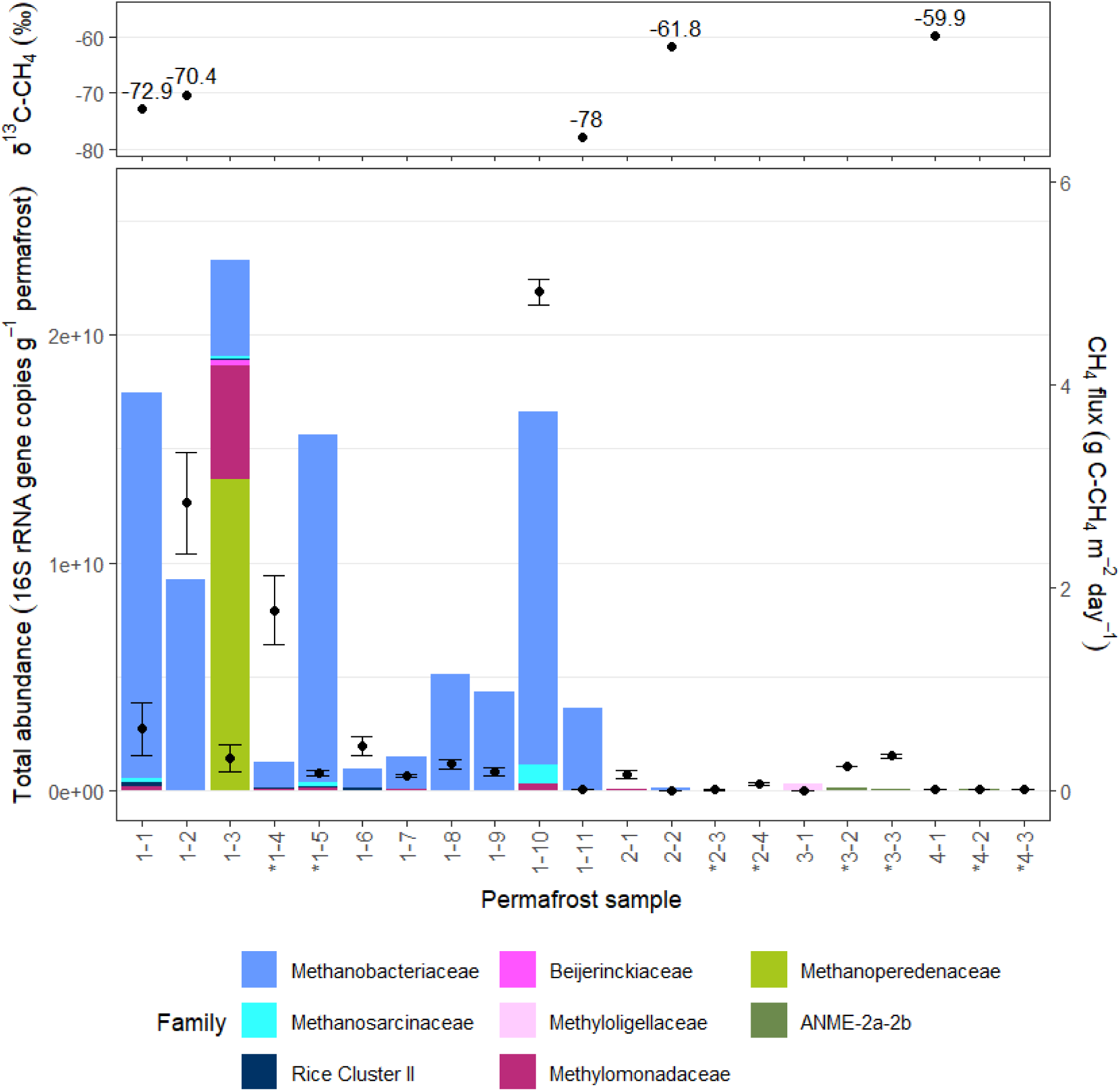
Methane isotopic ratios (top plot) compared to abundances of CH_4_ cycling bacteria and archaea (left axis) and CH_4_ fluxes (right axis). Abundances of methanogenic (blue), aerobic methanotrophic (pink), and anaerobic methanotrophic (ANME; green) taxa were identified for each wall chamber sample through 16S rRNA gene amplicon sequencing and scaled according to 16S rRNA gene copies determined through dPCR. Mean CH_4_ fluxes (right axis) is shown as a point for each permafrost sample (n = 3 replicates per sample). Samples are ordered from left to right by depth within in each permafrost unit. Winter samples are indicated by a star.

Permafrost-derived CH_4_ had low δ^13^C values consistent with microbial methanogenesis: -72.9 ‰ (32 ppm), -78.0 ‰ (36 ppm), and -70.4 ‰ (7 ppm) in Unit 1. While Units 2 and 4 had heavier, but still biogenic signatures of -61.8 ‰ (3 ppm) and -59.9 ‰ (7 ppm), they are still characterized as microbial in origin^38^. Samples depleted in δ^13^C-CH_4_ were enriched in δ^13^C-CO_2_, with values ranging from -15.8 to -11.0 ‰. This further supports biological methanogenesis as the source of emissions and indicates that CH_4_ was likely produced through hydrogenotrophy (CO_2_ reduction). Although we have no direct evidence of active methanotrophy in these permafrost samples (e.g. we see no CH_4_ uptake in our chamber measurements), *in-situ* CH_4_ oxidation is a possible cause for higher δ^13^C-CH_4_ values in Units 2 and 4. Ambient atmospheric gases collected above and upwind of the RTS had higher isotope values than those from the headwall: δ^13^C-CH_4_ was of -47.3 ‰ (2 ppm) and δ^13^C-CO_2_ was -14.1 ‰ (406 ppm), reflecting the regional influences from nearby thermogenic methane seeps^31^. A Keeling plot constructed from all CH_4_ chamber measurements (Fig. S1) yielded a y-intercept of -76 ‰ (R^2^ = 0.85), indicating that microbial methanogenesis is the source of the emitted CH_4_ ^38^. The heavier δ^13^C-CH_4_ values measured in Units 2 and 4 may be explained by dilution effects due to low CH_4_ concentrations, combined with mixing with atmospheric CH_4_.

To test the possibility that active microbial methanogenesis at sub-zero temperatures was the source of trapped gases, RNA was extracted from permafrost that was incubated for 24 hours at -5°C. All samples tested (1-1, 1-2, 1-4, 1-10, and 3-3), contained active cryophilic microbiota as indicated by detectable 16S rRNA. The *mcrA gene* which encodes for the alpha subunit of methyl-coenzyme M reductase, was actively transcribed in Unit 1 samples 1-1, 1-2, and 1-4, with transcript abundances ranging from 10^2^ to 10^3^ copies per g permafrost (6 x 10^3^ in 1-1, 3 x 10^2^ in 1-2, and 9 x 10^2^ in 1-4). No *mcrA* transcripts were detected in samples 1-10 and 3-3, which had the highest CH_4_ fluxes of their respective units. Nonetheless, these results demonstrate transcription of a key gene in the methane production pathway by active microbiota in permafrost, consistent with a microbial community active at sub-zero temperatures producing gases that may become trapped within the permafrost matrix.

### UAV methane mapping corroborates localized emissions

Lowlllaltitude uncrewed aerial vehicle (UAV) in-situ CH_4_ surveys detected enhancements directly next to the exposed headwall relative to background air (Fig. S2). Passes within ∼15 m of the upper thaw face showed elevated CH_4_ concentrations up to 2271 ppb against a background with a mean of 2065 ± SD 29 ppb. Concentrations near the base of the headwall approached background (down to ∼2030 ppb); however, no definitive measurable distinction could be made between exposures of units 1-4, due to the combination of the rotor wash, ground friction-induced wind turbulence, and survey resolution. These observations independently corroborate chamber measurements and demonstrate localized methane emissions from the RTS headwall as measured by a non-contact method.

### Distribution of methanogenic and methanotrophic microbiota

The community composition of CH_4_ cycling taxa differed significantly between cryostratigraphic units (PERMANOVA, Bray-Curtis, *R^2^* = 0.24, *p* = 0). TOC and TN also influenced community composition (Fig. S3); however, they were not included in the PERMANOVA test due to strong collinearity with cryostratigraphy, TOC, and TN (Table S5). There was no correlation between absolute abundances of methanogens or methanotrophs with GHG emissions in individual permafrost samples (Table S6).

Methanogenic archaea were abundant in Unit 1 permafrost, which had the highest methane emissions (10^2^ to 10^6^ 16S rRNA gene copies g^-1^ permafrost). Methanogenic microbiota were not diverse, with a single genus (*Methanobacterium*) as the dominant taxon across all samples, representing ∼98% of all identified methanogenic archaea. Only 3 families were identified at the site, belonging to *Methanobacteriaceae* (3 amplicon sequence variants; ASVs), *Methanosarcinaceae* (1 ASV), and Rice Cluster II (1 ASV). Mirroring heterogeneity in observed methane flux, *Methanobacterium* (and all methane generating taxa) were present in variable amounts representing <0.01 to ∼14% of microbial community composition (Fig. 4).

Methanogenic taxa were found in low relative abundance in Unit 2 (<0.3%) and were absent in Units 3 and 4. All methanogenic taxa identified at this RTS were capable of hydrogenotrophic methanogenesis (Fig. 5), a pathway that reduces CO_2_ and utilizes H_2_ as the primary electron donor. This is consistent with isotopically light CH_4_ and isotopically heavy CO_2_ from emitted gases, indicating that CH_4_ was produced through reduction of CO_2_. Although the presence of hydrogenotrophic taxa suggested a reliance on availability of CO_2_, there was no statistical relationship (Table S6) between CO_2_ flux and total abundance of microbiota (PERMANOVA, Bray-Curtis, R^2^ = 0.08, *p* = 0.13) or distribution of CH_4_ cycling taxa (Spearman’s *rho* = 0.18, *p* = 0.27).

**Fig 5.**
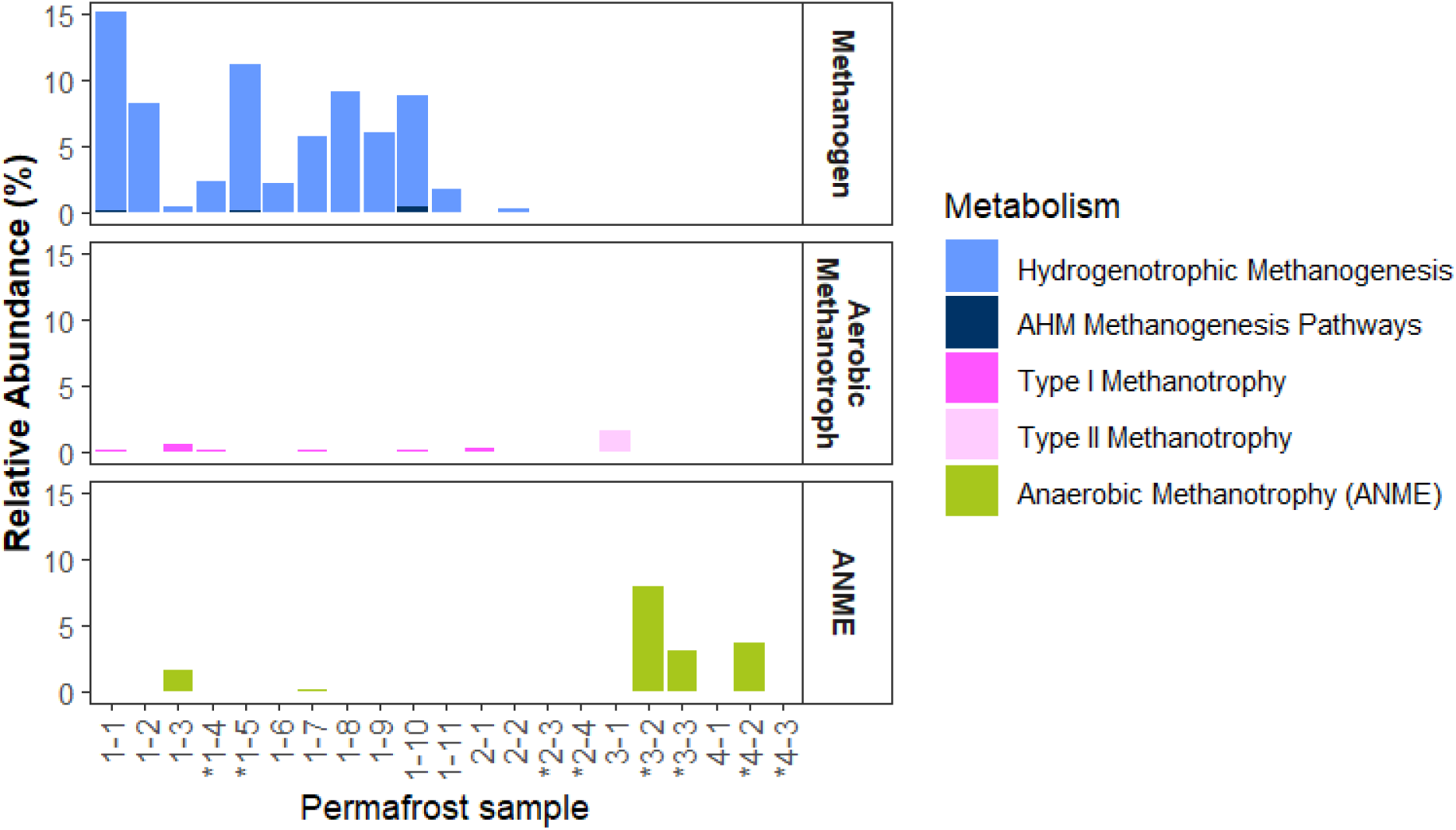
Relative abundances of CH_4_ cycling metabolisms from each permafrost core corresponding to in-situ wall chambers, based on taxa identified through 16S rRNA gene amplicon sequencing. Taxa capable of methanogenesis, aerobic CH_4_ oxidation, and anaerobic CH_4_ oxidation (ANME) are identified by shades of blue, pink, and green, respectively. Samples are ordered from left to right by depth within in each permafrost unit. The metabolism “AHM” indicates a taxon (Methanosarcinaceaea) that is capable of three methanogenesis pathways, including acetoclastic, hydrogenotrophic, and methylotrophic methanogenesis. Winter samples are indicated by a star

Microbiota that can oxidize CH_4_ were found in all cryostratigraphic units, though the dominant CH_4_ oxidizing pathway differed across units (Fig. 5). Type I aerobic CH_4_ oxidizing bacteria (*Methylomonadaceae*) were present in Unit 1 permafrost, most notably in samples with the highest CH_4_ fluxes (eg. Fig. 4, samples 1-4 and 1-10). Type II methanotrophs (*Beijerinckiaceae* and *Methyloligellaceae*) were present in low relative abundance (<3%) in samples with low CH_4_ flux (1-3 and 3-1). Anaerobic methane-oxidizing (ANME) archaea were identified across all permafrost units, though *Methanoperedenaceae* occurred only in Units 1 and 2 and ANME-2a-2b was limited to Units 3 and 4. For most ANME archaea, anaerobic oxidation of methane is only energetically favourable when a syntrophic partner acts as an external electron acceptor during reverse methanogenesis ^39^. Archaea in the ANME-2 clade live in consortia with sulfate reducing bacteria (SRB) and couple anaerobic CH_4_ oxidation with sulfate reduction^40^. Seven SRB genera were identified across all permafrost samples (Fig. S4) and were in higher abundance where ANME-2a-2b was present, indicating that anaerobic CH_4_ oxidation was possible in all permafrost samples. Several ANME archaea, including ANME-2, are capable of methylotrophic methanogenesis when CH_4_ concentrations are low and SRB have outcompeted methanogenic archaea for substrates^39,41^. ANME-2a-2b was only identified in Units 3 and 4, where no other methanogens are present and methane availability was low.

### Annual emissions from Niglintgak Island RTS

Between 2019 and 2024, this RTS ablated ∼90 m on the West side at the local topographic high and ∼50 m on the East side where there is a hillslope that is orthogonal to the RTS thaw face (Fig. S5 and S6), resulting in an erosion rate of 10 to 18 m per year (Fig. S7). In 2023 the retreating headwall was 205 linear m with an average height of 6.03 m. The surface area of exposed permafrost at the headwall was 1237 m^2^ (Fig. S5, Table S7).

Annual GHG emissions were estimated by summing median CH_4_ and CO_2_ fluxes from each permafrost unit measured during the summer (ie. when permafrost ablation occurs) and scaling to the surface areas of each cryostratigraphic unit. We estimate that this RTS on Niglintgak Island annually emits ∼9 kg C-CH_4_ and 100 kg C-CO_2_ immediately upon permafrost ablation at the headwall, resulting in emissions of ∼300 kg C as CO_2_ equivalents (CO_2_e) annually.

These emissions are independent of any secondary emissions from microbial decomposition of thawed carbon in the debris flow. Methane only comprised ∼7% of annual carbon emissions directly from the thaw face, however that proportion increased to 68% once converted to CO_2_e. This highlights the significant warming potential of carbon emissions released immediately during headwall ablation at RTS that are widespread throughout the Arctic.

## Discussion

This study provides the first direct measurements of GHG fluxes from permafrost exposed at the headwall of an active RTS. We show that CH_4_ and CO_2_ are immediately released to the atmosphere during permafrost thaw due to headwall retreat, revealing a previously unquantified natural GHG emission source in rapidly warming Arctic landscapes. Our observations indicate the source of GHG emissions from the headwall can be attributed to direct release of GHGs occluded in the permafrost matrix rather than from secondary mineralization stimulated by increased temperatures. Although trapped gases have been documented in permafrost in a variety of settings^6–11^, the origins of these gases have only been hypothesized. Two mechanisms have been suggested: (1) gases produced at warm temperatures (>0°C) could become trapped upon initial permafrost aggradation^42^, or (2) slow GHG production by cold-adapted microbiota that can accumulate within permafrost over millennia^6,9^. Our results support the latter. Cold-adapted methanogenic archaea actively transcribed the *mcrA* gene in samples incubated at -5°C, demonstrating that methanogenesis is possible under sub-zero conditions in these soils. Following the Holocene Thermal Maximum in this region (∼11.6 to 10.3ka), permafrost now exposed by the RTS gradually cooled as summer air temperatures transitioned to near-modern day temperatures between 6.7 and 5.6ka^43^, and since the late 1960s and early 1970s mean annual near-surface permafrost temperatures at this site have increase by ∼2 °C to the present range of ∼-6 to -7°C^44^, therefore active *in-situ* methanogenesis is possible in permafrost prior to RTS initiation. Cryophilic microbiota capable of activity and cell division at sub-zero temperatures have been found in permafrost soils globally nearly ubiquitously^45^.

Consistent with the interpretation of active cold-adapted microbiota as the primary CH_4_ source, is the presence of methanogenic archaea in Unit 1 permafrost, which is the cryostratigraphic unit with the highest methane fluxes, whereas the other units with lower CH_4_ emissions had few (<1% relative abundance) to no methanogens present. Isotopic data further demonstrates the microbial origin of the trapped CH_4_. Low δ^13^C-CH_4_ values (-78.0 to -70.4 ‰) and high δ^13^C-CO_2_ values (-14.4 to -11.0 ‰) in Unit 1 are characteristic of microbial hydrogenotrophic methanogenesis. The comparatively enriched (but still biogenic) δ^13^C-CH_4_ values in Unit 2 (-61.8 ‰) and Unit 4 (-59.9 ‰) could result from mixing of low CH_4_ values with background atmospheric CH_4_, or oxidation of CH_4_ by methanotrophs. These lines of evidence do not exclude the possibility that some GHGs could have been trapped during permafrost aggradation, but the data taken together show that biogenic CH_4_ is stored and subsequently released upon permafrost ablation, and that the permafrost contains active microbiota transcribing the key gene for methanogenesis.

Past research on permafrost carbon emissions has focused on the context of gradual active-layer deepening, with more recent work highlighting the importance of abrupt thaw processes that result in a complete change of state (eg. RTS, coastal and river bank erosion, and active layer detachment)^18,19^. Rapid thaw emissions have now been incorporated into new carbon models^19,46–48^, however, these models only consider carbon stocks and post-thaw mineralisation, not the immediate degassing of trapped CH_4_ and CO_2_. Current projections reflect large uncertainties in carbon that could be released due to rapid permafrost thaw, ranging from ∼0.28 PgC per year (80 ± 19 PgC by 2300) from all rapid thaw features^19^, to ∼1.95 x 10^3^ PgC per year from RTSs alone^48^. These estimates omit the contribution of immediate emissions from exposed headwalls. Our findings further demonstrate a GHG source that has never been included in permafrost-carbon feedback models, therefore even models that do account for rapid thaw features likely underestimate their emissions.

To date, there are limited prior studies (n=3) that directly measure GHG flux associated with RTSs; however, these studies measure emissions from the thawed debris-flow material of the RTSs^27–29^ and only one of these studies measured CH_4_ flux^27^. We measured headwall CO_2_ fluxes ranging from 1.4 x 10^-3^ to 6.1 g C-CO_2_ m^-2^ day^-1^, which is more variable but overlaps with observations at an RTS outflow in Siberia (0.24 to 2.6 g C-CO_2_ m^-2^ day^-1^)^27^ and RTS outflows in Alaska (2.07 ± 0.2 g C m^-2^ day^-1^)^28^. The CO_2_ flux values measured at the Niglintgak RTS are orders of magnitude lower than modeled fluxes based on mineralisation rates from laboratory permafrost incubations (∼367 g C m^-2^ year^-1^)^46^, though the emission rates measured here should be considered in addition to previous estimates of carbon flux. CO_2_ fluxes did not correlate with any measured physicochemical properties, indicating that CO_2_ emissions may be influenced by processes other than bulk abiotic characteristics of the permafrost.

CH_4_ fluxes measured here ranged from 4.8 x 10^-4^ to 5.0 g C-CH_4_ m^-2^ day^-1^. Although wetlands remain the largest natural global CH_4_ source (80-280 Tg yr^-1^)^49^, mean CH_4_ fluxes from the Niglintgak RTS (6.3 x 10^-1^ g C-CH_4_ m^-2^ day^-1^) were one order of magnitude higher than mean wetland fluxes in the High, Low, and Sub- Arctic^50^. The mean CH_4_ emission rates measured here were also one order of magnitude higher than northern peatland ponds, glacial and post-glacial lakes, and thermokarst lakes, but were similar to those in northern beaver ponds^51^. We estimate that direct release of CH_4_ emissions from this single RTS could reach ∼9 kg C-CH_4_ annually. Once converted into CO_2_e, release of trapped CH_4_ constituted 68% of radiative forcing from carbon emissions at this site. This is much higher than previous estimates which suggest CH_4_ comprises 50% of radiative forcing from rapid permafrost thaw^19^.

Immediate trapped gas release from permafrost exposed at RTS headwalls could be a significant natural source of GHG emissions in the Arctic, especially as the frequency and severity of RTSs are increasing with climate change. There were 2747 medium to large RTSs and 2244 to 12,514 small RTSs identified across the Northern Hemisphere between 2012 and 2022^48^, and thousands of new RTSs initiate across the Arctic every year^22,24–26^. We estimate that approximately 300 kg CO_2_e could be released annually from the headwall of the RTS on Niglintgak Island. Additionally, permafrost headwalls are produced by other abrupt processes, including coastal and riverbank erosion and active-layer detachment^18^, and thus represent an additional source of pan-Arctic GHG emissions for directly exposed permafrost. The pronounced heterogeneity of GHG fluxes, soil chemistry, and distribution of CH_4_ cycling taxa across different cryostratigraphic layers within a single RTS highlights the challenges in scaling site-level observations to the landscape or pan-Arctic scale. Broader field studies on RTS across the Arctic are needed to quantify this emerging component of the permafrost-carbon feedbacks to climate change.

### Conclusions

Current estimations of carbon emissions based on carbon stocks, laboratory microbial mineralization rates, and field studies on thawed material do not account for CH_4_ and CO_2_ emissions from immediate release of trapped gases from thawing permafrost headwalls. Direct field measurements of GHG fluxes from rapid thaw features are rare in the literature, and we show here that immediate release of trapped gases from permafrost exposures may be significant and is currently unaccounted for in climate models. Considering the quantity of current and future active RTS across the Arctic and ongoing erosion of permafrost coasts, including the direct release of trapped GHGs from exposed permafrost headwalls in climate models could significantly lower uncertainties of permafrost carbon feedbacks on climate.

## Methods

### Radiocarbon dating

Sub-samples of permafrost cores collected from each cryostratigraphic unit were thawed and, together with active layer samples, oven dried (105°C) for 24 hours. Pieces of relatively intact organic detritus including roots were hand-picked from the samples under a microscope, and remaining bulk samples were ground to a fine powder (Table S2). Accelerator mass spectrometry (AMS) ^14^C ages were determined for the bulk soil samples and organic detritus to help establish a general site history. Dating was completed by the A.E. Lalonde AMS Laboratory (Ottawa, Canada). Calibration was performed using OxCal v4.4^52^ and the IntCal20 calibration curve^53^. Confidence intervals for calibrated ages (cal BP) reported here are within 2σ.

### In-situ CH_4_ and CO_2_ flux measurements

Shallow (10–34 cm) horizontal boreholes were drilled into the frozen headwall using a 2.375-inch O.D. (6.03 cm) frozen soil coring barrel auger (AMS, American Falls, ID) driven by a battery powered hammer drill. Sample locations were selected based on visibly distinct cryostratigraphic permafrost layers.

Soil gas flux chambers were constructed from an ABS pipe (5 cm inner diameter, 6 cm outer diameter), outfitted with a removable ABS threaded plug with o-ring seal and two luer lock fittings to allow continuous air flow to the gas analyser when sealed. Open soil chambers were inserted into each borehole and allowed to freeze in place for ∼1 hr before applying the plug. The chambers were a fixed length (∼10 cm) whereas borehole length varied; therefore, the exposed permafrost area differed among measurements. This area was accounted for in the surface area calculation for each measurement. It was not possible to install the chambers 24 hours prior to measurements during summer because ongoing thaw and retreat of the RTS headwall caused the chambers to become unstable. For consistency, the same 1-hour timing was therefore used during winter measurements.

For each chamber, CH_4_ and CO_2_ concentrations were measured over 30 to 300 s observation lengths (including dead band) using a Licor 7810 (LI-COR Biosciences, Lincoln, USA) portable GHG analyzer (1 Hz measurement rate, 250 standard cubic centimeters per minute flow rate). Measurements were repeated in triplicate for each borehole with the chambers open between measurements, allowing CH_4_ and CO_2_ concentrations to reach background ambient levels (∼2 ppm CH_4_; ∼400 ppm CO_2_) prior to the next measurement. Flux measurements were taken on August 15, 2022, April 20, 2023, and August 10, 2023.

Retrieved cores were placed in sterile Whirl-Pak bags and placed into a cooler with ice packs for ∼4 hours until transported back to the Western Arctic Research Centre, Inuvik, where we stored them in a –20°C freezer until they were transported to the lab in Guelph, ON. The cores remained frozen for the duration of travel.

### Near-surface atmospheric CH_4_ mapping

Near-surface CH_4_ concentrations were mapped using an UAV, a DJI Matrice 300 RTK equipped with a prototype open-path mid-infrared tunable diode laser absorption spectrometer (TDLAS). This instrument has a sensitivity of 8 ppb and a 10 Hz sampling rate, making it capable of resolving low-concentration enhancements emitted from low-flux sources. Further information regarding this sensor design can be found in Beattie *et al*. (currently under review)^54^.

Flight operations were conducted on August 10, 2023. The UAV was manually flown in a quasil11grid pattern across the RTS and up the headwall face at ∼0.5–25 m above or, in the case of the headwall, away from the surface. Background transects were additionally flown upwind from the RTS. To limit rotorl11induced mixing of the air and influence the measurements, we maintained low airspeeds (up to 3.5 m/s) with a lateral standoff from the face and with repeated passes. CH_4_ time series were synchronized with RTK/GNSS positions and mapped in QGIS 3.34.

### Carbon isotope geochemistry

Gas samples were collected for carbon isotope ratio analysis from each borehole (1-1, 1-2, 1-11, 2-2, 3-1, and 4-1) on August 15, 2022. Following each flux measurement, the chambers remained sealed to permit gas accumulation, and discrete gas samples (∼100 mL) were collected from each unit (except Unit 3, which was lost in transit) and stored in 1 L foil bags (Restek, #22950). A discrete air sample (∼100 mL into a 1 L foil bag) was also collected on nearby undisturbed tundra, upwind from the RTS, to serve as a background for the measurements. Samples were measured for carbon isotope ratios of CH_4_ and CO_2_ via cavity ring down spectroscopy (G2201i, Picarro, Santa Clara, USA) within two days of collection.

Calibrated standards were used for the full concentration range measured. Precision was up to 7 ‰ at the low air values, and above 3 ppm, precision was 2 ‰.

### Gas flux calculations

Concentrations of CH_4_ and CO_2_ were plotted against time for each replicate chamber measurement and the change in gas concentration over time was determined using the linear portion of each slope. Gas flux for each chamber measurement was calculated using the following formula ^55^,

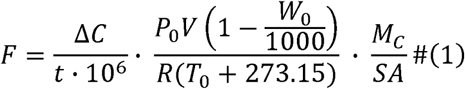

where *F* is the GHG flux (g C-(CH_4_ or CO_2_) m^-2^ day^-1^), Δ*C* is change in gas concentration (CO_2_ was measured in ppm; CH_4_ was measured in ppb and divided by 10^3^), *t* is time (s), *P_0_*is initial atmospheric pressure (atm), *V* is chamber volume (L), *W_0_*is the initial water vapor mole fraction (mmol mol^-1^) *R* is the ideal gas constant (L atm K^−1^ mol^−1^), *T_0_* is initial air temperature (°C), *M_C_* is molar mass of carbon (g/mol), and S*A* is surface area (m^2^) of exposed permafrost at the back of the wall chambers. Daily temperature and atmospheric pressure data for the observation period were obtained from Environment and Climate Change Canada Historical Climate Data (https://climate.weather.gc.ca/index_e.html) for the Pelly Island, NT weather station, located 31 km north of Niglintgak RTS.

### Trapped Gas Measurements

Trapped gases were measured as in Lapham *et al*., 2020^8^. In a walk-in freezer, frozen subsamples (∼5 g, 1 cm diameter) were collected, transferred to a vial, and sealed with a butyl rubber stopper. Each subsample was then rapidly thawed at room temperature, 8 mL of supersaturated brine was added, and the slurry was vortexed for 15 s. Gases were sampled from the headspace using a 20 mL syringe (20 mL removed from ∼60 mL headspace) and injected into a 12 mL evacuated glass exetainer (Labco, Lampeter, Wales, UK). Concentrations of CH_4_ and CO_2_ were measured using a gas chromatograph (Bruker 450 GC, Bruker Corporation, Billerica, USA).

### Mapping retreat rate and surface area of the slump

The rate of permafrost ablation at the active headwall was estimated by comparing satellite images from Sept 3, 2019 and July 21, 2024 (Google Earth Pro v 7.3.6.10201; Fig. S7). To quantify the surface area of the exposed permafrost headwall and individual cryostratigraphic units within the RTS, the RTS was surveyed on August 10, 2023 using a DJI Matrice 300 RTK UAV equipped with a DJI P1 camera. A high-overlap gridded flight captured detailed overhead images, followed by a free flight to obtain oblique views of the RTS face. The images captured by the UAV were later processed using structure-from-motion in Pix4DMapper (v3.2.23) to generate a high-resolution 3D model (Fig. S5). Within the software, a 3D multifaceted surface object was created for each permafrost unit to delineate polygonal areas of exposed permafrost, and the calculated areas (Table S7) were subsequently used in emission estimates.

### Estimation of annual emissions

Annual CH_4_ and CO_2_ emissions were estimated based on the summer median GHG fluxes for each permafrost unit. It was assumed that exposure of permafrost and release of trapped GHGs at the thaw face of the RTS headwall would primarily occur between June and September (122 days), when the average air temperature is above 0°C on Niglintgak Island. During this time period, the unfrozen water content of permafrost behind the thaw face increases according to soil freezing characteristics as ground temperature approaches 0°C^56^, and permafrost ablation occurs at the headwall^20^. In the winter months, limited permafrost thaw at south-facing RTS headwalls can occur despite sub-freezing air temperatures due to microclimatic effects^57^, however it was assumed that unfrozen water contents were predominantly near zero and permafrost ablation was halted. Therefore, winter GHG fluxes were attributed to trapped gases released by drilling and core excavation rather than from natural permafrost thaw. Annual GHG emissions were calculated by summing median CH_4_ and CO_2_ fluxes from each permafrost unit measured during the summer (ie. thaw season), using the following formula

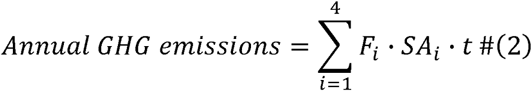

where *i* is the permafrost unit, *F* is median GHG fluxes (g C-(CH_4_ or CO_2_) m^-2^ day^-1^) from summer measurements (Table S8), *SA* is the surface area of the exposed permafrost unit (m^2^; Table S7), and *t* is number of days with average air temperature above 0°C (122 days).

CO_2_ equivalents (CO_2_e) were calculated by multiplying annual CH_4_ emissions by a factor of 27^58^ and summing with annual CO_2_ emissions.

### DNA extractions and 16S rRNA gene sequencing

The outer ∼2 mm of each frozen (stored at -20°C) permafrost core was scraped off with sterile tools to remove potential contaminants introduced during drilling. DNA was extracted from each core alongside a negative control using the DNeasy Powerlyzer Powersoil kit (Qiagen, Hilden, Germany), according to the manufacturer’s protocol with minor modifications: homogenization was completed by 45 s bead beating three times; eluent (sterile water) was incubated on the spin column membrane for five minutes at room temperature; elution step was carried out twice with 25 μL nuclease-free H_2_O (total 50 μL DNA extract).

DNA extracts were used for amplicon sequencing of the 16S rRNA gene at the Integrated Microbiome Resource at Dalhousie University (Halifax, Canada) using universal V4-V5 primers 515F 5’- GTGYCAGCMGCCGCGGTAA-3’ and 926R 5’-CCGYCAATTYMTTTRAGTTT-3’^59^ on an Illumina MiSeq in a paired end (300 bp PE) sequencing run as described in ^60^. The genomic data generated for this study are available in the NCBI Genbank database under BioProject PRJNA1088168. Sample names used in this study and the coordinating sample names submitted to NCBI can be found in Table S9.

Amplicon Sequence Variants (ASVs) were identified using the DADA2 pipeline v 1.26.0^61^ in R v 4.2.0^62^. Sequences were truncated at 290 bp for forward reads and 210 bp for reverse reads and had primers removed. Sequences were assigned taxonomy using the Silva database v 138.1^63^. Sample data (GHG fluxes, physicochemical soil properties, and 16S rRNA gene copies) was integrated into microbial assemblage data using Phyloseq v 1.42.0^64^. Absolute abundances of taxa were determined by multiplying relative abundances by the mean quantity of 16S rRNA gene copies g^-1^ permafrost.

Prior to statistical analysis, the taxonomic data was subset to only include methanogenic and methanotrophic taxa. The vegan v 2.6-4^65^ PERMANOVA (adonis2) function was used to statistically determine if variables had a significant effect on microbial distribution based on the Bray-Curtis dissimilarity matrix. Principal coordinates analysis (PcoA) was carried out using the Bray-Curtis dissimilarity matrix to visually represent groupings in the dataset. Scripts used for bioinformatic analyses were deposited into a github repository (https://github.com/lexojoyo/permafrost_amplicon).

### Permafrost incubation, RNA extraction, and cDNA synthesis

To determine if microbial communities could be active in permafrost at sub-zero temperatures, permafrost cores with high CH_4_ flux and/or high abundances of methanogens respective to its permafrost unit (1-1, 1-2, 1-4, 1-10, 3-3) were sub-sampled and incubated for 24 h at -5°C. RNA was extracted from each core alongside a negative control using the RNeasy PowerSoil Total RNA Kit (Qiagen), according to manufacturer protocol using phenol:chloroform:isoamyl alcohol (25:24:1, v/v; pH 6.7) with one modification: elution was carried out with 50 µL molecular H_2_O to concentrate the final extract.

RNA extracts were treated with DNase I (New England Biolabs, Ipswich, USA), using 50 µL of RNA extract, 6 µL of DNase I reaction buffer, 2 µL of DNase I, and 2 µL of molecular H_2_O. Removal of genomic DNA was confirmed by the absence of visible bands on an agarose gel after targeting the V4-V5 region of the 16S rRNA gene with PCR. First-strand complementary DNA (cDNA) was synthesized using the SuperScript IV First-Strand Synthesis System (Invitrogen, Waltham, USA). The reverse transcription assays were 20 µL and contained 5 µL RNA extract, 1 µL random primers, 1 µL 10 mM dNTP mix, 4 µL 5X SSIV Buffer, 1 µL 100 mM DTT, 1 µL Ribonuclease Inhibitor, 1 µL SuperScript IV Reverse Transcriptase, and 6 µL RNase-and nuclease-free H_2_O. Reverse transcription was performed by incubating at 23°C for 10 min and 50°C for 10 min. Reverse transcriptase was inactivated by incubating at 80°C for 10 min.

### Quantification of the 16S rRNA gene

The 16S rRNA gene was quantified in DNA extracts from permafrost and cDNA libraries from laboratory incubations using quantitative PCR (qPCR) to determine bacterial and archaeal cell abundance. Assays were run on the Quantstudio 3 Real-Time PCR System (Thermo Fisher Scientific, Waltham, USA). Each qPCR assay was 20 μL and contained 10 μL PowerTrack SYBR Green Master Mix (Thermo Fisher Scientific), 6.4 μL molecular H_2_O, 2 μL of template (diluted 1:10), and 0.8 μL each of 10 μM forward primer 515F (5’-GTGCCAGCMGCCGCGGTAA-3’) and reverse primer 806R (5’-GGACTACHVGGGTWTCTAAT-3’)^66^. Purified DNA from a bacterial isolate was used as a positive control, and molecular H_2_O was used in place of a template for the non-template control. All sample and control assays were run in triplicate. The qPCR cycling conditions were 95°C for 2 min, followed by 40 cycles of denaturation at 95°C for 15 s and annealing and extension at 52°C for 1 min. Double-stranded gBlock Gene Fragments (Integrated DNA Technologies, Coralville, USA) spanning the V4-V5 region of the 16S rRNA gene were used to produce a standard curve (R^2^ = 0.992, efficiency = 98.42).

### Quantification of the *mcrA* gene

The presence of the *mcrA* gene in cDNA was assessed by PCR amplification. Samples that yielded visual bands with agarose gel electrophoresis (1-1, 1-2, and 1-4) were selected to be quantified with digital PCR (dPCR).

To determine the quantity of *mcrA* genes transcribed in incubated permafrost samples, digital PCR (dPCR) was performed on cDNA using using the Thermo Fisher Scientific Quantstudio Absolute Q Digital PCR System.

A 10 μL mastermix solution was prepared with 2 μL Absolute Q Universal DNA dPCR Master Mix (Thermo Fisher Scientific), 0.2 μL 25X SYBR Green I gel stain, 5.4 μL molecular H_2_O, 2 μL of template cDNA, and 0.2 μL each of 10 μM forward primer mlas-F (5’-GGTGGTGTM GGDTTCACMCARTA-3’) and reverse primer mcrA-rev (5’-CGTTCATBGCGTA GTTVGGRTAGT-3’)^67^. Nine μL of mastermix solution was loaded into individual wells in a Quantstudio Absolute Q MAP 16 plate (Thermo Fisher Scientific) and overlain with 15 μL Quantstudio Absolute Q Isolation Buffer (Thermo Fisher Scientific). The dPCR cycling conditions were 95°C for 10 min, followed by 40 cycles at 95°C for 15 s and 50°C for 1 min. Assays were analyzed using the Quantstudio Absolute Q Digital PCR Software (v6.3.2). The threshold was positioned above the negative partitions (4000 relative fluorescence units), using the positive and negative control as a reference. Accepted assays had distinguishable separation between the positive and negative partitions, and >20,000 partitions that passed the ROX quality control. A sample was deemed positive if ≥ 1 partition was detected.

### Physicochemical soil properties

Prior to measuring carbon and nitrogen content, subsamples of each permafrost core were air dried at room temperature and ground using a ball mill (Retsch PM100, Haan, Germany). Total carbon (TC) was analyzed by dry combustion (*ca*. 250 mg) at 1300°C^68^ using a Leco carbon analyzer (CR-12, St. Joseph, USA). A second soil subsample (*ca.* 5 g) was ignited at 475°C for four hours in a muffle furnace to remove organic matter and was subsequently measured (*ca.* 250 mg) for C content by dry combustion as above^69^; this second subsample represented total inorganic carbon (TIC). Total organic carbon (TOC) was determined by subtracting TIC from TC. Total nitrogen (TN) was measured on *ca.* 250 mg tin-encapsulated samples by combustion (i.e., Dumas method) using a Leco FP-528 nitrogen determinator with an oven temperature of 900°C and reduction column set to 750°C^70^.

Bulk density (g/ml) of the permafrost cores was determined by dividing the mass of each frozen core by its volume. We determined dry-basis gravimetric water content (*w*) of each permafrost core according to the following equations,

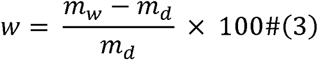

Where *m_w_*is the mass (g) of the wet sample and *m_d_* is the stable mass (g) reached after samples were air dried at room temperature.

### Statistical analyses

Statistical analyses were performed using R v 4.2.0^62^. All GHG flux and physicochemical data failed tests for normality using the Shapiro-Wilk test. Data transformations did not improve data distribution and boxplots indicated heteroskedasticity in the variance across groups (cryostratigraphic units), therefore the Kruskal-Wallis test and Dunn’s one-sided post-hoc test was used to determine statistical differences between variables in each permafrost unit. As this data did not fit assumptions of normality, homogeneity, or linearity, Spearman’s rank-order correlation was used to assess correlation between GHG fluxes and physicochemical soil data. The *p* values were adjusted using the Holm-Bonferroni method for both Dunn’s test and Spearman’s correlation.

## Acknowledgements

We are grateful to Claudia Wood, Elisse Magnuson, Brendan O’Neill and Wendy E. Sladen for field work assistance, Maureen Strauss for measuring stable isotopes, and Jenna Scharnowski for assistance with molecular work. Our field work was supported by local wildlife monitors Miles Dillon, Christine Firth, and Todd Gruben. Fieldwork was conducted under Northwest Territories Scientific Research Licence nos. 17074 and 17466. This research was conducted on current and ancestral territories of many Indigenous and First Nations peoples, including the Inuvialuit and Nihtat Gwich’in Nation (Tuktoyaktuk and Inuvik, NWT); the ancestral lands of the Attiwonderonk and the Haudenosaunee people, and the territory of the Mississaugas of the Credit First Nation (Guelph, ON); the unceded territory of the Anishinàbe Algonquin People (Ottawa, ON); the traditional territory of the WLJSÁNEĆ (Pauquachin, Tsartlip, Tsawout, Tseycum) peoples and the Malahat First Nation (Sidney, BC); and the Piscataway People (Solomons, MD).

