## Supplementary Materials for "Immediate methane and carbon dioxide release from exposed permafrost at an active retrogressive thaw slump in the Canadian Arctic"

**Supplementary Information**

**1. Supplemental Methods**

**Measurement of trapped gases**

Although trapped gases were detected in all samples, it is unclear whether the measured quantities accurately represent the *in-situ* concentration of trapped gases due to the potential for gas loss prior to extraction. The destructive nature of permafrost drilling certainly disrupts and releases gases that were previously trapped, at least along the surface area of the permafrost that was in contact with the core barrel. The small diameter of cores (5 cm) also promotes diffusion of gases to the exterior of the core during transport; while cores remained largely frozen, the exterior of cores were “soft” and for the first few hours in the field, cores were stored in a cooler with ice packs and warmed significantly from in situ temperatures. Additionally, previous work has shown that trapped CH_4_ can be released from permafrost at temperatures as low as -20°C^1^, therefore it is possible that gases may have been lost during storage as well. We cannot draw any conclusions regarding the distribution of trapped gases throughout the geologic units due to these uncertainties, however it is clear that measurable quantities of CH_4_ and CO_2_ are trapped within the frozen permafrost matrix and are released upon thaw. Future work to measure this should use larger diameter cores, or place cores in supersaturated brines immediately upon collection.

**2. Supplemental Results**

**Geological context and permafrost unit interpretations**

The Niglintgak RTS is retreating into ice-rich permafrost within a morainal deposit emplaced during the Early Wisconsinan Toker Point Stade (Buckland Glaciation)^2^. Following permafrost establishment, the terrain of this region was modified by thaw during the early Holocene warm interval and the active layer increased to about 2.5 times thicker than at present, creating a regionally expressed thaw unconformity dated to ~8000 ^14^C years BP (~9000 calendar years BP)^3^. Throughout the Holocene, solifluction and soil creep have been continuously redistributing materials downslope^2^, promoting formation of a buried organic layer at slope bases^4^.

Unit 1 (cryoturbated organic-rich slope deposits) is underlying the 30–50 cm modern active layer with earth hummocks, restricted to the southern/eastern flank where the thaw face intersects a slope. Radiocarbon dates from bulk sediment and organic detritus range from 6180 – 4420 calBP (5260 ± 20 to 4020 ± 20 ^14^C years BP). This unit has variable ice content with thin ice lenses and veins. The heterogeneous mixture of dark, organic-rich and medium-brown sandy silt reflects reworking by cryoturbation, soil creep, and solifluction. Organic material accumulates between hummocks and at the base of the active layer due to cryoturbation and is progressively frozen as the surface and permafrost aggrades, forming a buried organic layer^4,5^.

Unit 2 (paleoactive-layer formed during the Holocene Thermal Maximum) is medium-brown sandy silt, with visible roots. There is variable ice content in thin lenses and veins, and some reticulate ice cryostructures that often indicate freezing of saturated sediments^4,5^. A sharp thaw unconformity separates Unit 2 from 3. Hand-picked organic materials yielded an age of 9400 - 9030 calBP (8240 ± 25 ^14^C years BP) and bulk soil yielded an age ranging from 30720 – 30030 calBP (26100 ± 110 to 25900 ± 110 ^14^C years BP). The relatively young age of hand-picked organic materials from Unit 2 suggests that the bulk sediments would likely have an age more similar to basal ice (Unit 4) and refrozen melt-out till (Unit 3), but the average age is younger due to the influence of this added organic material. Regionally, Unit 2 represents a paleo-active layer ~2.5 x thicker than present developed during the Holocene Thermal Maximum. This unit is widespread in this region and is associated with a basal thaw unconformity dating from approximately 8000 ^14^C years BP^3^.

Unit 3 (refrozen saturated sediments) is a medium-grey silty diamicton above a thaw unconformity with underlying Unit 4. It contains reticulate ice cryostructures indicative of slow freezing of saturated sediments. This unit lacks the visible roots, which is congruent with underlying Unit 4. Radiocarbon ages of Unit 3 (40 380 – 39 070 calBP; 34400 ± 290 ^14^C years BP) indicate close temporal proximity to Unit 4, and reflect thaw, saturation and refreezing of supra-glacial melt-out till.

Unit 4 (basal glacial icy sediments) is located at the base of the exposure and is consistent with glacially deformed icy sediments described by Murton *et al*., (2005)^6^. It comprised of a medium-grey silt rich diamicton and massive ice in deformed layers. Radiocarbon dating yielded an age of 41 440 – 40 080 calBP (35 800 ± 340 ^14^C years BP), which is consistent with the non-finite date (>37 000 ^14^C BP) reported for the same locality (Site No. 9 in Rampton, 1988). Pieces of coal and rare wood are evident in the RTS debris flow, and organic material, likely coal, was visible in permafrost samples under a microscope. We do not attribute the age of this unit to a specific geocryologial event because of inclusion of older wood and charcoal that may be substantially older than the deposition event; however, the materials predate the Late Wisconsinan advance of the Laurentide Ice Sheet that began 27 000 – 30 000 ^14^C years BP and overrode the region by about 23 000 – 24 000 ^14^C years BP ^7^, during the Toker Point Stade (Rampton, 1988). We interpret Unit 4 to be basal ice of the Laurentide Ice Sheet overlain by Unit 3 that, following Murton *et al.* (2005), we subsequently interpret as supraglacial melt-out till formed from thaw of higher portions of the basal icy layer and subsequent preservation by permafrost aggradation.

**Microbial ecology was influenced by permafrost physicochemical properties**

Bacterial and archaeal community structure was influenced by geologic unit (PERMANOVA, Bray-Curtis, *p* < 0.001). Unit 1 permafrost was the most diverse, and microbial diversity decreased in older permafrost layers. The higher concentrations of essential nutrients such as TOC and TN could likely support a wider range of microorganisms that may not be well adapted to low nutrient conditions. Thus, the taxonomic diversity of Unit 1 permafrost was more consistent with modern soils, including common soil microbiota such as *Acidobacteriae, Actinobaceteria, Bacteroidota,* and *Clostridia* (Fig. S4). Total organic carbon, total nitrogen, and gravimetric water content were significantly lower in Units 2 through 4, shifting the microbial community to taxa that were better adapted to such oligotrophic conditions. Unit 2 permafrost was less diverse than Unit 1 and the most abundant taxon was a soil bacterium (*Gemmatimonadaceae*) known for its ability to adapt to low water availability^8^. *Clostridium* sensu stricto *spp*. were the most abundant taxa identified in all permafrost layers except for Unit 2, where they were notably absent from three of the four samples sequenced. Microbiota appeared to be distinctly adapted to the extreme environmental conditions in Units 3 and 4. Of the few taxa identified in these oldest permafrost layers, most were anaerobic, endospore-forming, or were commonly found in cold and extreme environments. *Clostridium* sensu strico *spp*. constituted up to 73% relative abundance in individual samples in Units 3 and 4. This genus forms endospores under environmental stress, therefore their high abundance indicates dormancy as a likely survival strategy in these cold and oligotrophic soils.Caldatribacteriota

Sequencing of the 16S rRNA gene yielded a total of 119,675 reads with 1222 ASVs identified from all 21 samples. Low quantities of reads (10^2^ – 10^4^) were recovered from each sample. Few sequence reads may indicate low microbial biomass in the soil, however low sequence recovery could also be a consequence of environmental substances (eg. humic acids) inhibiting DNA extractions and PCR (a common issue in permafrost soils), or technical issues during the sequencing run. This makes results difficult to interpret, as the true microbial diversity and abundance may be higher than what is presented here. To determine if low read recovery was due to low cell concentrations in the samples, qPCR was performed to quantify 16S rRNA gene copies as a proxy for community biomass. Microbial biomass was variable across samples, ranging from 1.4 x 10^9^ to 8.0 x 10^11^.16S rRNA gene copies per g permafrost (Table S4). The bacterial and archaeal biomass as measured by qPCR was highly correlated with number of recovered reads per permafrost sample (Spearman’s *rho* = 0.56, *p* < 0.001). Microbial biomass was significantly higher in Unit 1 permafrost compared to the older, less organic permafrost units (Kruskal-Wallis and Dunn, *p* < 0.01).

**3. Supplemental Figures**

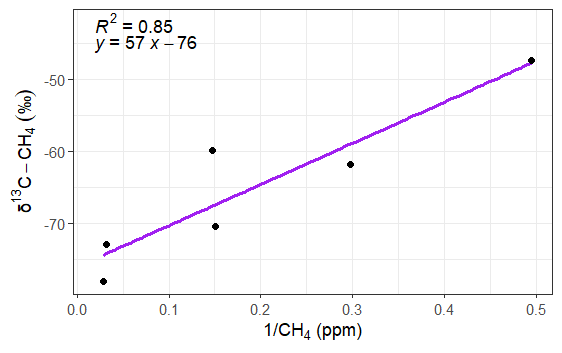

**Fig S1.** Keeling plot of δ^13^C-CH_4_ values and CH_4_ concentration.

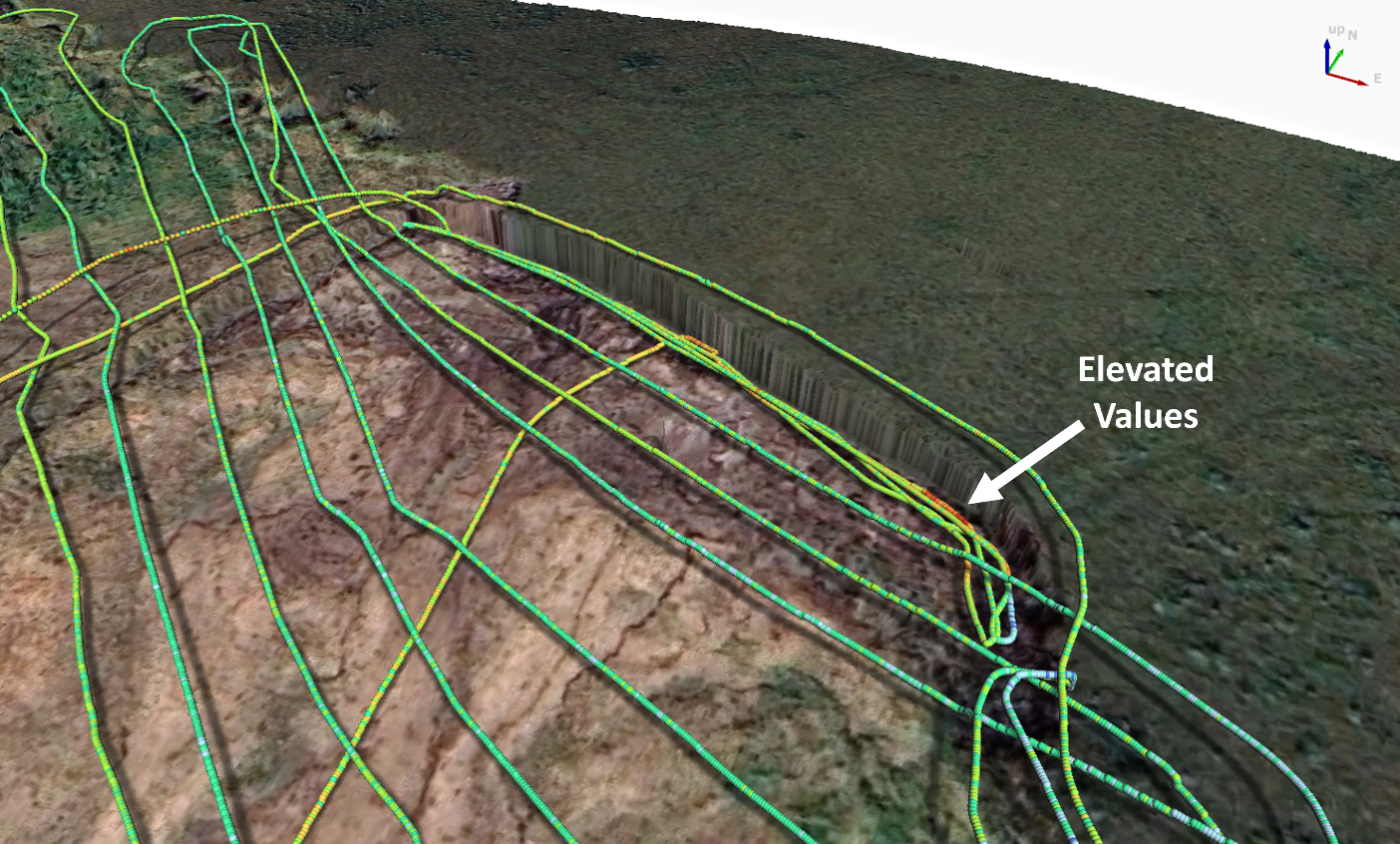

**Fig S2.** Near‑surface UAV methane mapping over the RTS from August 10, 2023. Flight tracks are draped on the SfM surface and colour‑coded by CH_4_ concentration. Near‑source passes along the upper headwall (within ~15 m of the face) show elevated values relative to background (mean: 2.065 ± SD: 0.029 ppm) of up to 2.271 ppm. Enhancements decay to background downwind of the face. Apparent differences at the same plan‑view locations reflect pass‑to‑pass changes in flight altitude/standoff and minute‑scale wind variability. The 3D rendering shows altitude along tracks to aid in interpretation.

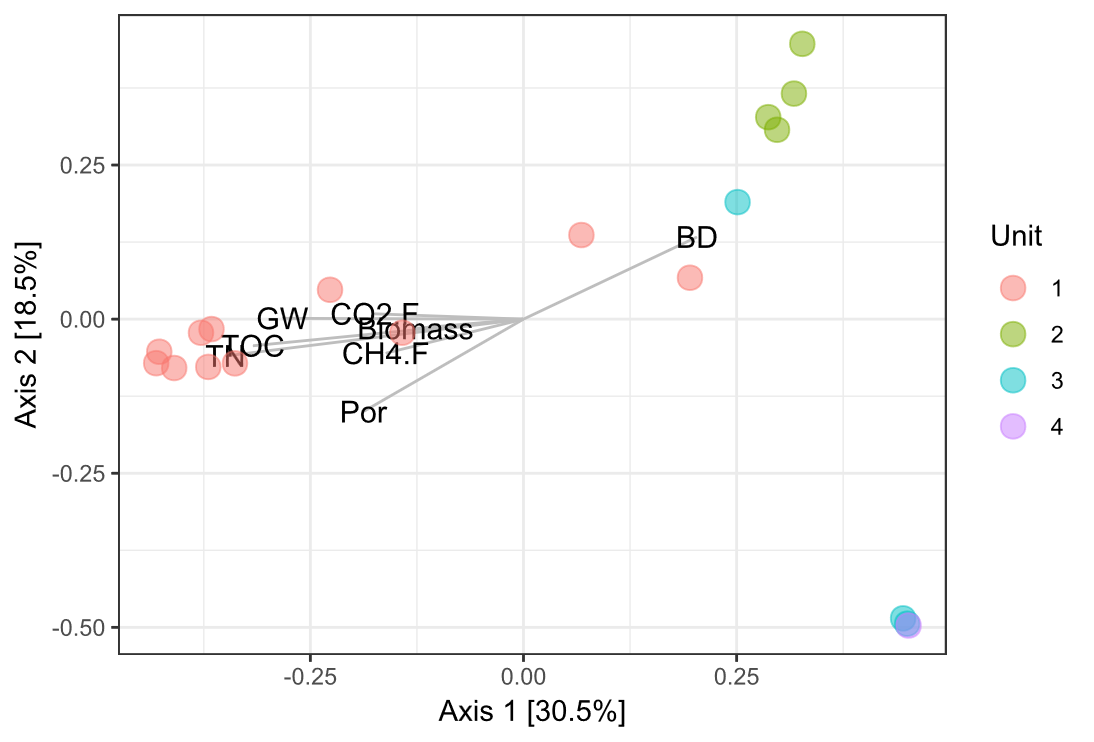

**Fig. S3.** Principal coordinates analysis (PcoA) of absolute abundances of methanogenic and methanotrophic ASVs from permafrost samples (*n* = 19). No CH_4_ cycling taxa were identified in samples 4-1 and 4-3, therefore they were excluded from this analysis. Bray-Curtis dissimilarity was used (PERMANOVA, *p* = 0.001) and permutations were set to 999. Axis 1 and Axis 2 accounted for 30.5 and 18.5%, respectively, of the differences in the methane cycling communities in these samples. Points are coloured according to cryostratigraphic units. Biomass represents 16S gene copies g^-1^ permafrost as measured through qPCR, CH4.F is CH_4_ flux, CO2.F is CO_2_ flux, TOC is total organic carbon, TN is total nitrogen, Por is porosity, GW is gravimetric water content, and BD is bulk density.

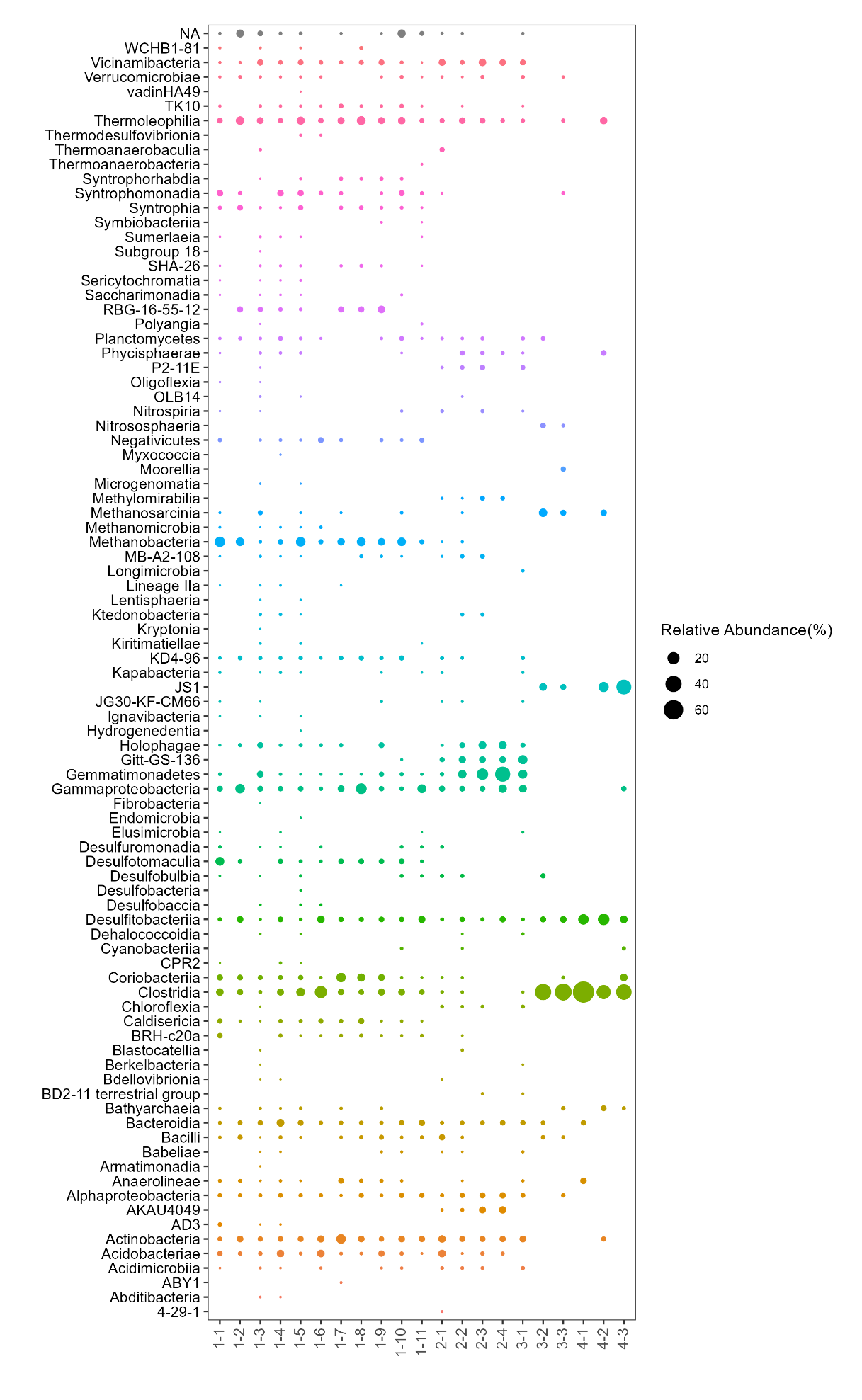

**Fig S4.** Relative abundances of all Bacterial and Archaeal classes (represented by different colours), identified through 16S rRNA gene amplicon sequencing of each permafrost core collected.

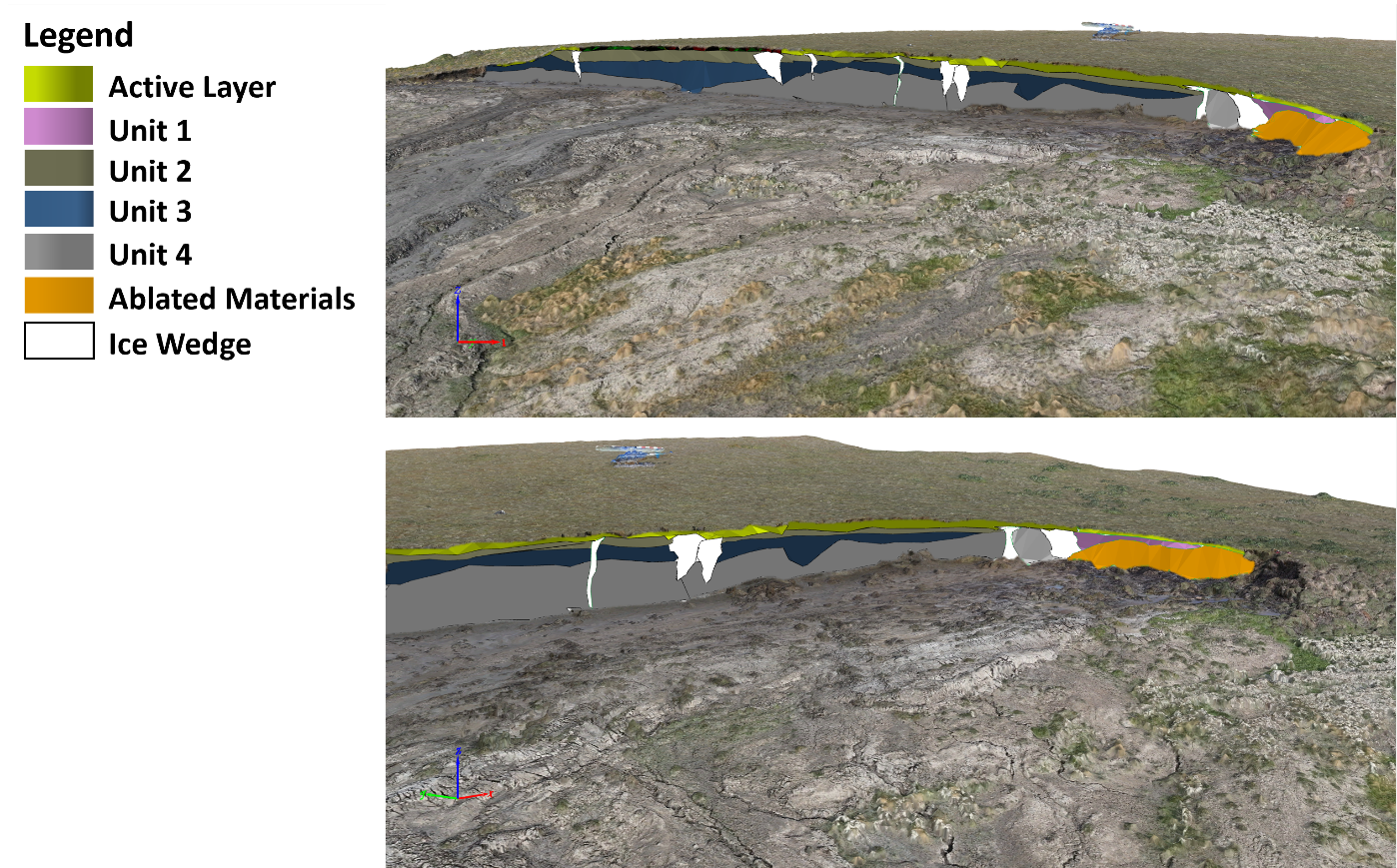

**Fig S5.** Structure‑from‑motion (SfM) 3D model of the RTS headwall at Niglintgak Island from UAV imagery acquired on August 10, 2023. Two oblique perspectives showing the exposed permafrost face with 3D delineations of cryostratigraphic units draped on the mesh; colours are the same as Fig. 2 and correspond to the active layer (green), Unit 2 (dark brown), Unit 3 (blue-grey), Unit 4 (grey), and ice wedge (light grey/white).

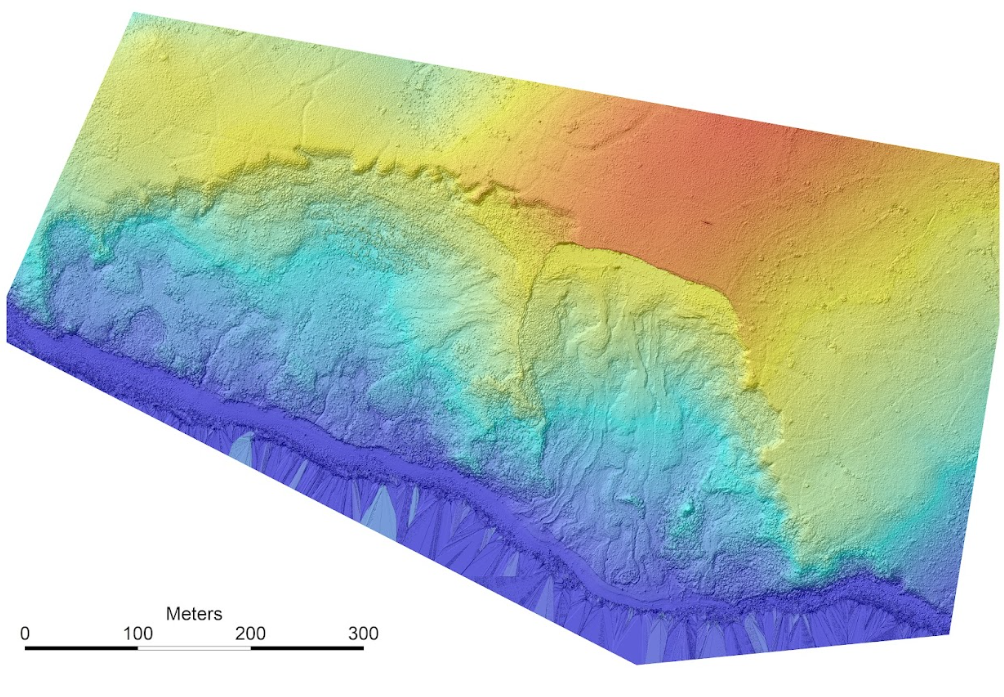

**Fig S6**. Structure-from-motion digital surface model of the RTS for August 10, 2023, created using photographs taken by a UAV.

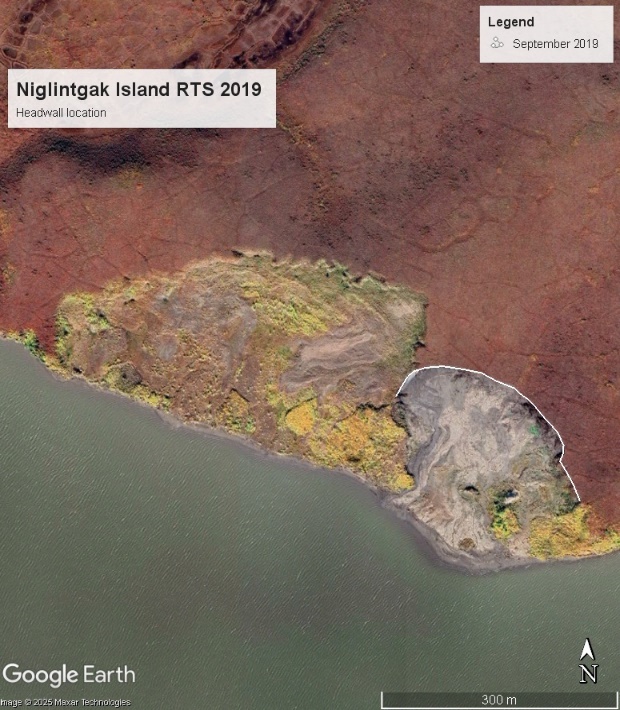

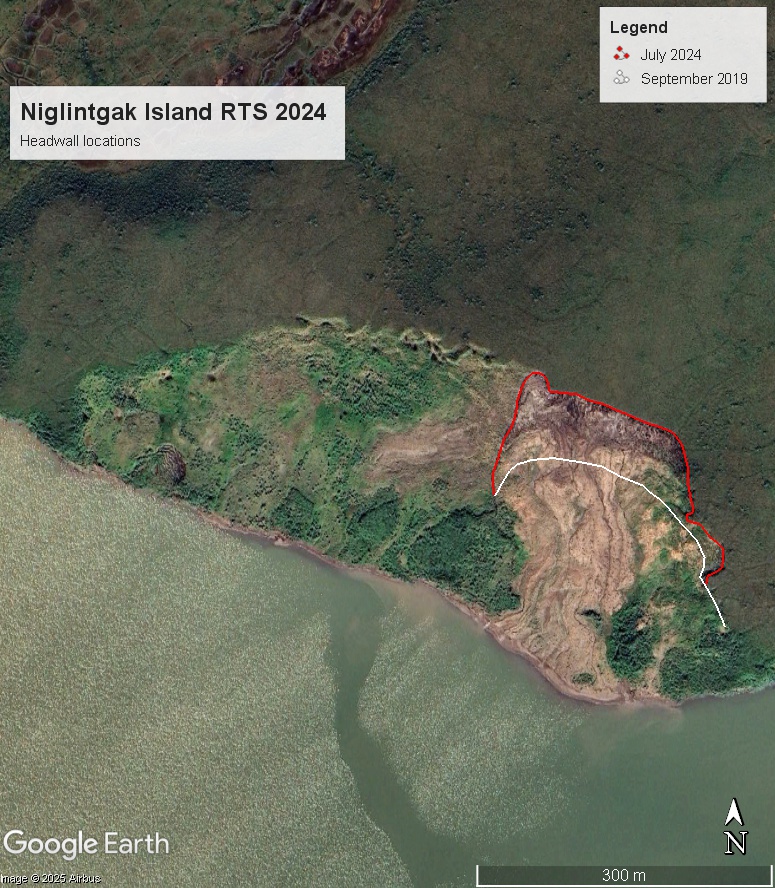

**Fig S7.** RTS headwall locations in 2019 (left) and 2024 (right).

**4. Supplemental Tables**

**Table S1.** Mean physicochemical properties, biomass, and total reads from each permafrost sample.

| **Sample** | **Trapped CH_4_ (mg C-CH_4_ g^-1^ permafrost)** | **Trapped CO_2_ (mg C-CO_2_ g^-1^ permafrost)** | **TOC (%)** | **TN (%)** | **Bulk density (g cm^-3^)** | **Gravimetric water (g g^-1^)** | **Biomass**  **(16S rRNA gene copies g^-1^ permafrost)** | | **Total reads** |
| --- | --- | --- | --- | --- | --- | --- | --- | --- | --- |
| 1-1 | 2.1E-02 | 1.6E-01 | 13.60 | 0.73 | 1.0 | 1.4 | | 1.13E+11 | 10 800 |
| 1-2 | 1.4E-02 | 1.9E-01 | 13.97 | 1.02 | 0.9 | 2.6 | | 1.11E+11 | 754*^w^* |
| 1-3 | 1.3E-02 | 2.8E-01 | 21.38 | 0.84 | 1.2 | 1.8 | | 8.00E+11 | 52 400 |
| 1-4 | 4.7E-02 | 5.5E-01 | 14.48 | 0.58 | 1.3 | 2.7 | | 4.61E+10 | 1 040 |
| 1-5 | 1.1E-01 | 7.2E-01 | 33.80 | 1.26 | 1.7 | 2.1 | | 1.37E+11 | 2 460*^w^* |
| 1-6 | 8.2E-02 | 5.7E-01 | 27.37 | 0.86 | 1.1 | 3.1 | | 4.47E+10 | 1 940*^w^* |
| 1-7 | 1.1E-02 | 2.1E-01 | 12.29 | 0.52 | 0.8 | 5.8 | | 2.49E+10 | 14 200*^w^* |
| 1-8 | 5.7E-02 | 5.8E-01 | 15.70 | 0.66 | 1.0 | 3.8 | | 5.52E+10 | 1 940*^w^* |
| 1-9 | 5.1E-02 | 5.2E-01 | 19.08 | 0.85 | 0.9 | 2.5 | | 7.20E+10 | 3 220*^w^* |
| 1-10 | 6.1E-03 | 2.3E-01 | 39.15 | 1.69 | 0.8 | 3.7 | | 1.86E+11 | 640*^w^* |
| 1-11 | 1.3E-02 | 2.1E-01 | 21.24 | 0.66 | 1.1 | 1.7 | | 2.09E+11 | 230 |
| 2-1 | 2.5E-03 | 2.0E-01 | 2.65 | 0.13 | 1.0 | 1.4 | | 1.49E+10 | 255*^w^* |
| 2-2 | 6.4E-04 | 2.2E-02 | 1.21 | 0.08 | 1.2 | 0.9 | | 2.60E+10 | 2 600 |
| 2-3 | 1.8E-04 | 3.3E-02 | 1.01 | 0.07 | 1.9 | 0.2 | | 7.36E+10 | 162*^w^* |
| 2-4 | 9.6E-05 | 4.3E-02 | 1.19 | 0.06 | 1.6 | 0.3 | | 3.34E+10 | 685*^w^* |
| 3-1 | 1.7E-06 | 1.4E-02 | 1.31 | 0.09 | 1.8 | 0.4 | | 1.89E+10 | 263*^w^* |
| 3-2 | 9.8E-04 | 2.4E-02 | 1.85 | 0.10 | 0.9 | 0.7 | | 1.40E+09 | 108*^w^* |
| 3-3 | 8.9E-04 | 2.6E-02 | 1.72 | 0.09 | 1.4 | 0.5 | | 2.51E+09 | 10 008*^w^* |
| 4-1 | 1.5E-04 | 4.4E-02 | 1.30 | 0.10 | 1.2 | 1.4 | | 4.59E+09 | 754*^w^* |
| 4-2 | 1.6E-03 | 3.4E-02 | 1.78 | 0.10 | 1.5 | 0.5 | | 1.93E+09 | 52 400*^w^* |
| 4-3 | 1.2E-03 | 3.1E-02 | 1.25 | 0.07 | 1.8 | 0.2 | | 2.70E+09 | 1 040*^w^* |

TOC: total organic carbon; TN: total nitrogen. *^w^* indicates samples that had “weak” sequencing runs, as identified by the sequencing centre (IMR, Dalhousie, NS).

**Table S2.** Radiocarbon dating results.

| Unit | Material | ^14^C yr BP | ± | F^14^C | ± | calBP |
| --- | --- | --- | --- | --- | --- | --- |
| 1 | soil | 5260 | 20 | 0.5196 | 0.0012 | 6180 - 5940 (95.4%) |
| 1 | soil | 4130 | 20 | 0.5981 | 0.0013 | 4820 - 4530 (95.4%) |
| 1 | organic detritus | 4910 | 20 | 0.5429 | 0.0012 | 5660 - 5590 (95.4%) |
| 1 | organic detritus | 4020 | 20 | 0.6068 | 0.0013 | 4520 - 4420 (95.4%) |
| 2 | soil | 26100 | 110 | 0.0391 | 0.0005 | 30720 - 30030 (95.4%) |
| 2 | soil | 25900 | 110 | 0.0399 | 0.0005 | 30310 - 29980 (95.4%) |
| 2 | organic detritus | 8240 | 25 | 0.3589 | 0.0010 | 9400 - 9030 (95.4%) |
| 3 | soil | 34400 | 290 | 0.0138 | 0.0005 | 40380 - 39070 (95.4%) |
| 4 | soil | 35800 | 340 | 0.0117 | 0.0005 | 41440 - 40080 (95.4%) |

Note: Errors for Conventional Radicocarbon Ages (^14^C yr BP) are 1σ. Confidence intervals for calibrated ages (calBP) are within 2σ. Calibration was performed using OxCal v4.4 ^9^ and the IntCal20 calibration curve ^10^.

**Table S3.** Minimum, maximum, median, and mean CH_4_ and CO_2_ fluxes for the RTS as a whole and for each permafrost unit (encompassing both seasons).

|  | **CH_4_ flux (g CH_4_-C m^-2^ day^-1^)** | | | | **CO_2_ flux (g CO_2_-C m^-2^ day^-1^)** | | | |
| --- | --- | --- | --- | --- | --- | --- | --- | --- |
|  | Min | Max | Median | Mean | Min | Max | Median | Mean |
| Entire RTS | 4.8 x 10^-4^ | 5.0 | 1.9 x 10^-1^ | 6.3 x 10^-1^ | 1.4 x 10^-3^ | 6.1 | 4.5 x 10^-1^ | 8.4 x 10^-1^ |
| Unit 1 | 1.9 x 10^-2^ | 5.0 | 3.2 x 10^-1^ | 1.1 | 1.4 x 10^-3^ | 6.1 | 5.7 x 10^-1^ | 1.2 |
| Unit 2 | 4.8 x 10^-4^ | 2.1 x 10^-1^ | 3.6 x 10^-2^ | 6.2 x 10^-2^ | 6.6 x 10^-2^ | 1.2 | 4.6 x 10^-1^ | 5.4 x 10^-1^ |
| Unit 3 | 6.2 x 10^-4^ | 3.7 x 10^-1^ | 2.4 x 10^-1^ | 2.0 x 10^-1^ | 4.0 x 10^-2^ | 3.4 x 10^-1^ | 1.1 x 10^-1^ | 1.7 x 10^-1^ |
| Unit 4 | 1.6 x 10^-2^ | 2.1 x 10^-2^ | 1.7 x 10^-2^ | 1.8 x 10^-2^ | 1.1 x 10^-2^ | 1.4 | 5.7 x 10^-1^ | 4.5 x 10^-1^ |

**Table S4.** Concentration and carbon isotope ratios from *in*-*situ* CH_4_ collected following flux wall chamber measurements.

| Permafrost sample | ^13^C-CH_4_ (‰) | CH_4_ concentration (ppm) | ^13^C-CO_2_ (‰) | CO_2_ concentration (ppm) |
| --- | --- | --- | --- | --- |
| 1-1 | -72.9 | 31.76 | -11.00 | 494.80 |
| 1-2 | -78.0 | 35.50 | -11.90 | 501.90 |
| 1-11 | -70.4 | 6.63 | -14.40 | 474.40 |
| 2-2 | -61.8 | 3.35 | -14.80 | 465.20 |
| 4-1 | -59.9 | 6.78 | -15.80 | 471.80 |
| Background atmospheric | -47.3 | 2.02 | -14.10 | 406.40 |

**Table S5.** Spearman *rho* correlation matrix for GHG flux and physicochemical soil properties.

|  | CO_2_ flux | CH_4_ flux | TOC | TN | *w* | ρ |
| --- | --- | --- | --- | --- | --- | --- |
| CO_2_ flux | 1.00 | 0.54* | 0.33 | 0.40 | 0.34 | -0.23 |
| CH_4_ flux |  | 1.00 | 0.65* | 0.68* | 0.62* | -0.54* |
| TOC |  |  | 1.00 | 0.94* | 0.80* | -0.51* |
| TN |  |  |  | 1.00 | 0.80* | -0.57* |
| *w* |  |  |  |  | 1.00 | -0.71* |
| ρ |  |  |  |  |  | 1.00 |

TOC: total organic carbon; TN: total nitrogen; *w*: gravimetric water; ρ: bulk density.
**p* < 0.01. The *p* values were adjusted using the Holm-Bonferroni method.

**Table S6.** Spearman *rho* correlation matrix for GHG flux, microbial total abundance, and CH_4_ cycling taxa.

|  | CO_2_ flux | CH_4_ flux | Total abundance | Methanogen | Methanotroph | ANME |
| --- | --- | --- | --- | --- | --- | --- |
| CO_2_ flux | 1 | 0.80^†^ | 0.36^†^ | 0.18^†^ | 0.07^†^ | -0.39^†^ |
| CH_4_ flux |  | 1 | 0.24^†^ | 0.23^†^ | -0.11^†^ | -0.19^†^ |
| Absolute abundance |  |  | 1 | 0.14^†^ | -0.06^†^ | -0.17^†^ |

^†^all *p* values > 0.01. The *p* values were adjusted using the Holm-Bonferroni method.

**Table S7.** Surface area of exposed permafrost at the RTS on Niglintgak Island. Surface areas (SA) were obtained using structure from motion mapping.

| Unit | SA (m^2^) | Mapped SA of exposed headwall permafrost (%) |
| --- | --- | --- |
| Unit 1 | 145.92 | 12 |
| Unit 2 | 264.59 | 21 |
| Unit 3 | 342.90 | 28 |
| Unit 4 | 483.98 | 39 |
| Total SA | 1237.39 | 100 |

**Table S8.** Summer (August) mean and median CH_4_ and CO_2_ fluxes for the RTS as a whole and for each permafrost unit. Median summer fluxes were used to calculate annual emissions from the RTS

|  | **CH_4_ flux (g CH_4_-C m^-2^ day^-1^)** | | **CO_2_ flux (g CO_2_-C m^-2^ day^-1^)** | |
| --- | --- | --- | --- | --- |
|  | Median | Mean | Median | Mean |
| Entire RTS | 2.1 x 10^-1^ | 7.8 x 10^-1^ | 5.5 x 10^-1^ | 1.1 |
| Unit 1 | 3.0 x 10^-1^ | 1.1 | 5.8 x 10^-1^ | 1.3 |
| Unit 2 | 6.2 x 10^-2^ | 8.4 x 10^-2^ | 3.5 x 10^-1^ | 3.5 x 10^-1^ |
| Unit 3 | 7.6 x 10^-4^ | 7.4 x 10^-4^ | 3.2 x 10^-1^ | 3.1 x 10^-1^ |
| Unit 4 | 1.9 x 10^-2^ | 1.9 x 10^-2^ | 1.26 | 1.26 |

**Table S9.** Sample names used in this study and the coordinating sample names submitted to NCBI Genbank database under BioProject PRJNA1088168 (Microbial characterisation of permafrost from the Inuvialuit Settlement region).

| Sample ID in this study | NCBI Genbank sample name | Permafrost unit | Date sampled |
| --- | --- | --- | --- |
| 1-1 | Nig22-1 | 1 | 2022-08-15 |
| 1-2 | Nig22-2 | 1 | 2022-08-15 |
| 1-3 | Nig23-Aug-1 | 1 | 2023-08-10 |
| 1-4 | Nig23-Apr-7 | 1 | 2023-04-20 |
| 1-5 | Nig23-Apr-8 | 1 | 2023-04-20 |
| 1-6 | Nig23-Aug-3 | 1 | 2023-08-10 |
| 1-7 | Nig23-Aug-10 | 1 | 2023-08-10 |
| 1-8 | Nig23-Aug-5 | 1 | 2023-08-10 |
| 1-9 | Nig23-Aug-8 | 1 | 2023-08-10 |
| 1-10 | Nig23-Aug-6 | 1 | 2023-08-10 |
| 1-11 | Nig22-3 | 1 | 2022-08-15 |
| 2-1 | Nig23-Aug-11 | 2 | 2023-08-10 |
| 2-2 | Nig22-4 | 2 | 2022-08-15 |
| 2-3 | Nig23-Apr-5 | 2 | 2023-04-20 |
| 2-4 | Nig23-Apr-6 | 2 | 2023-04-20 |
| 3-1 | Nig22-5 | 3 | 2022-08-15 |
| 3-2 | Nig23-Apr-3 | 3 | 2023-04-20 |
| 3-3 | Nig23-Apr-4 | 3 | 2023-04-20 |
| 4-1 | Nig22-6 | 4 | 2022-08-15 |
| 4-2 | Nig23-Apr-1 | 4 | 2023-04-20 |
| 4-3 | Nig23-Apr-2 | 4 | 2023-04-20 |
